# Replication fork directionality reveals how structural variants arise under replication stress

**DOI:** 10.64898/2026.06.29.735381

**Authors:** Dominik Glodzik, Mathilda Rigby, Michail Andreopoulos, Jake Crawford, Shannon Ehmsen, Avraam Tapinos, Alex Cornish, Richard Houlston, David C. Wedge, Ralph Scully, Peter J. Park

## Abstract

Structural variants (SVs) in cancer are associated with defects in DNA repair and replication stress, but the mechanisms generating common SV types remain unresolved. We propose that large (>100 kb) tandem duplications originate through a novel sister-fork breakage–fusion mechanism. To capture replication-related context beyond breakpoints, we developed an algorithm to characterize replication timing, origin density, and fork direction across SV-spanned regions, features that refine and differentiate previously defined SV signatures. Large tandem duplications frequently overlap replication origins from which forks proceed bidirectionally; combined with independent evidence from APOBEC strand asymmetry, this pattern is compatible uniquely with the proposed mechanism. Although tandem duplications in *CCNE1*-amplified and *CDK12*-mutant cancers also concentrate around origins and highly transcribed genes, they display distinct contexts: *CDK12*-mutant SVs arise near later-firing origins, whereas those in *CCNE1*-amplified tumors often coincide with genes in specific strand configurations, suggesting different causes of fork stalling. Incorporating replication features into signature analysis enabled the discovery of new SV signatures, which we used to build SVIG, a multi-class classifier of SV phenotypes. SV signatures attributed to replication stress may help guide therapies targeting this vulnerability.

## INTRODUCTION

Structural variants (SVs) in cancer are typically interpreted through their immediate consequences, such as oncogene amplification, tumor suppressor loss, or gene fusion. Equally important, deletion, tandem duplication, inversion, and translocation patterns reflect the cell’s DNA repair capacity and replicative state. For example, characteristic SV configurations are hallmarks of homologous recombination deficiency (HRD), a key determinant of therapeutic response.^1–3^ Recognizing these patterns helps identify patients likely to benefit from therapies exploiting synthetic lethality, such as PARP inhibitors.^4^

SV–based mutational signatures^5,6^ capture recurrent patterns across cancer cohorts. These signatures are derived from SV classifications by size (inter-breakpoint distance) and break-point type, distinguishing clustered (compound) events from singletons. Per-sample event counts are decomposed using non-negative matrix factorization or topic modelling^7^ to identify population-level patterns. Resulting signatures, e.g. RS1–20,^5^ are typically dominated by specific SV types or size ranges and have been linked to catastrophic chromosomal events^8^, DNA repair defects, or replication stress.^5^

Signatures of replication stress may refine patient stratification and improve clinical trial outcomes for emerging therapies targeting replication-associated vulnerabilities.^9–11^ For example, excess of tandem duplications (TDs) occurs in tumors with *CDK12* loss or *CCNE1* amplification.^9^ *CDK12* inactivation occurs in (5–6%) of prostate and (3–4%) of ovarian cancers^10,12^ and is associated with aggressive disease and poor prognosis.^13–15^ Similarly, *CCNE1* amplification occurs in roughly 20% of ovarian cancers, which are often resistant to platinum therapy.^16–18^ Despite their clinical significance and potential to indicate drug targets, the mechanisms underlying chromosomal instability in these patient populations remain unclear.

One reason why these mechanisms have not yet been resolved might be that current SV signatures do not incorporate key genomic features linked to SV formation, including replication dynamics (Figure 1A). For example, short (*∼*10 kb) TDs, which are abundant in *BRCA1* mutant cancers, have been linked to aberrant fork restart ( Figure 1B),^19^ or to microhomology-mediated break-induced replication (MMBIR, Figure 1C).^20^ Larger (>100 kb) tandem duplications, which are common throughout the cancer genomes, likely arise by mechanisms distinct from *BRCA1*-linked TDs. Motivated by observations of tandem duplications arising in yeast after re-replication,^21^ we recently proposed the sister-fork breakage–fusion (SFBF) mechanism,^3^ a hypothesis untested in human data. Aberrant fork restart involves converging replication tracts; in MMBIR, replication proceeds unidirectionally; and the SFBF model predicts that tandem duplications arise when two diverging replication forks, initiated bidirectionally from a single origin, stall and fuse (Figure 1D). The measurable abundance of replication origins and replication fork direction at SV loci can discriminate among candidate SV mechanisms and refine their signatures (Figure 1D).

**Figure 1.**
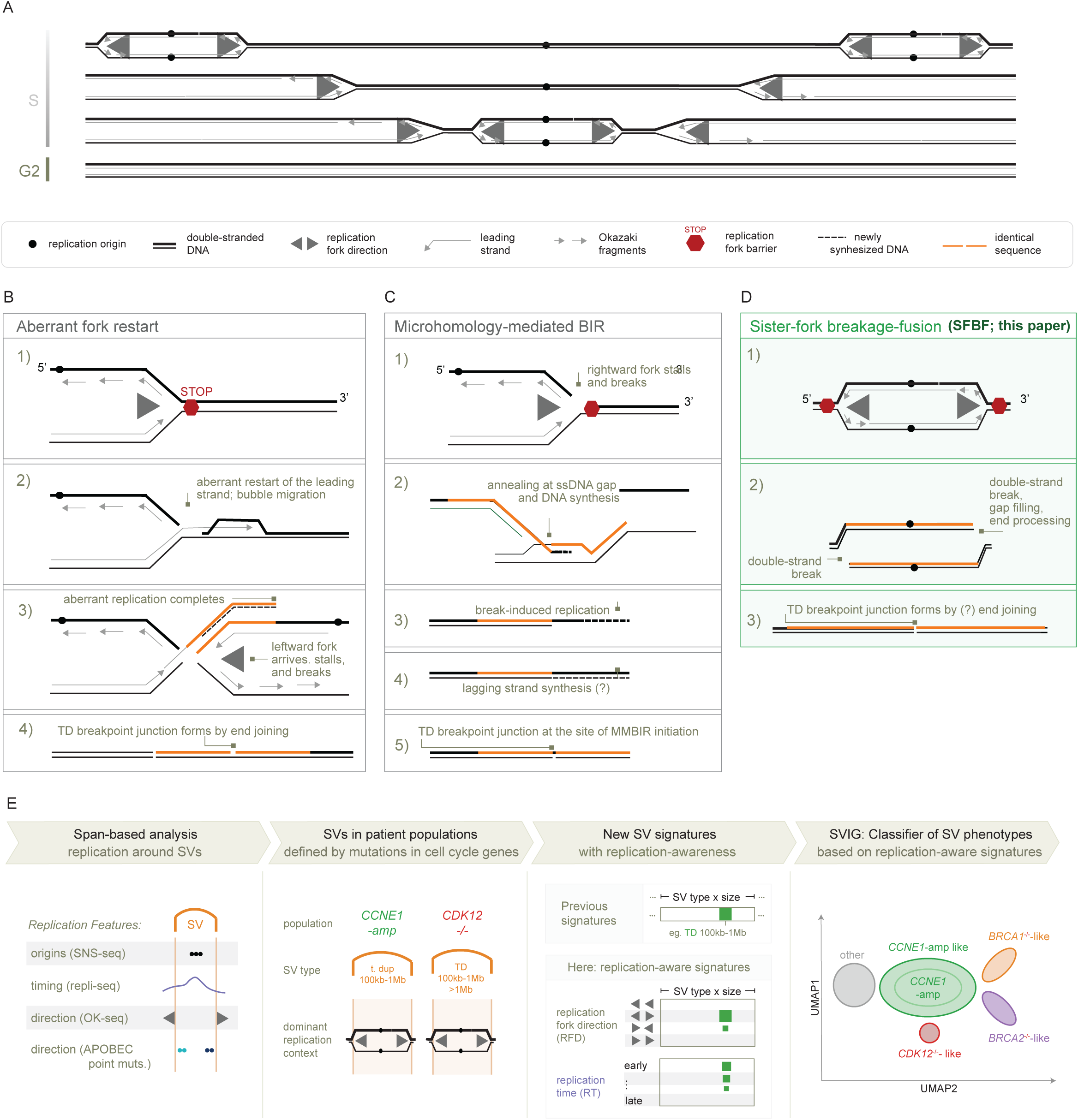
Models of tandem duplication (TD) formation following replication fork stalling. (A) Normal DNA replication, with replication forks firing throughout the S phase, does not result in SVs. (B) Aberrant fork restart. Converging or stalled forks undergo aberrant restart events that yield a TD. (C) Microhomology-mediated break-induced replication (MMBIR). A collapsed fork initiates a strand invasion and, consecutively, a TD. (D) Sister-fork breakage–fusion (SFBF) model. Two bi-directional forks initiated from a common origin stall and fuse, generating a TD. (E) Schematic and summary of replication-associated features at SV loci that are interrogated in this manuscript.

Here, we systematically analyze the relationship between SV signatures and local genomic context, including DNA replication and transcription. Unlike previous studies that focused solely on breakpoints (Figure S1),^6,22^ our algorithm examines the full genomic segments spanned by SVs and integrate co-occurring somatic point mutations to link SVs to replication dynamics. We then perform SV signature discovery using these features using the PCAWG dataset^23^. Finally, we develop a multi-class classifier of SV phenotypes and apply it to the Genomics England dataset—the largest dataset of SVs in cancer to date. Collectively, we demonstrate that SV features can directly capture replication stress in tumors.

## RESULTS

### Replication origins and fork direction shape SV positioning

To elucidate the mechanisms of structural variant (SV) formation, we systematically characterized replication features at loci spanned by SVs. The analysis focused on non-clustered SVs, as defined previously,^24^ to study variants arising independently of one another. We defined a normalized coordinate system centered at the SV midpoint (0), with the lower (5’) and upper (3’) breakpoint positions in the reference genome mapped to –0.5 and +0.5, respectively (Figure 2A). We reasoned that aligning replication profiles to standardized SV coordinates would enable visualization and comparison across SVs of varying lengths, summarizing replication origin density and timing at midpoints and breakpoints. Such aggregate replication profiles can discriminate between models of SV formation; for example, only the SFBF model predicts an excess of replication origins and earlier replication timing at SV midpoints relative to breakpoints (Figure 2B).^3^ We used SNS-seq–defined core replication origins, predicted to initiate 80% of forks across tested cell lines,^25^ together with MCF7 replication timing profiles from ENCODE (Repli-seq), selected as the closest match to the tissues of interest.^26,27^ To analyze specific SV classes, we assigned PCAWG SVs to previously defined rearrangement signatures (RS1–20; e.g., RS1).^5^ Each SV was assigned to the signature with the highest posterior probability, without applying a threshold, based on sample-level signature exposures and SV features, including type and size (inter-breakpoint distance).

**Figure 2.**
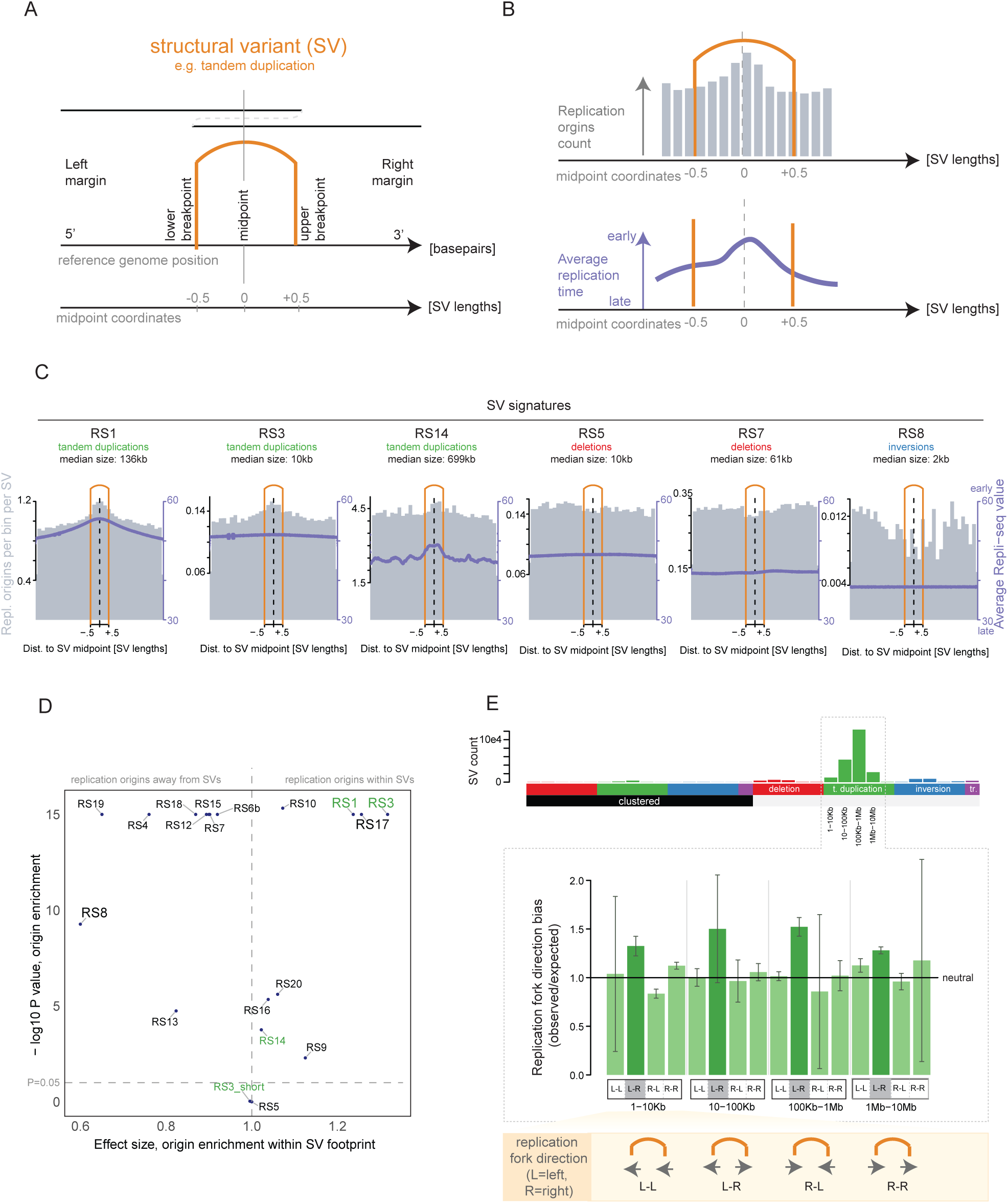
Segment-based analysis of SV loci in relation to replication timing and fork direction. (A) Schematic of an SV represented as two breakpoints mapped to the reference genome. To compare SVs of varying size, breakpoint pairs are transformed into a normalized coordinate system in which the SV midpoint is set to zero and the inter-breakpoint distance is scaled to one. (B) Expected placement of replication origins and associated replication timing profiles at tandem-duplication loci under the SFBF model. (C) Density of replication origins and average replication timing around SV breakpoints (orange) for common non-clustered SV signatures. RS1, RS3, and RS14 show profiles consistent with the SFBF model. Each flanking region spans three SV lengths. (D) Enrichment of replication origins within SV-spanning regions across SV signatures. Shown is a two-sided proportion test comparing the number of replication origins inside SVs versus within 1 Mb flanking regions. Tandem-duplication–dominated signatures (RS1, RS3) show the strongest enrichment, which disappears when restricting RS3 to SVs shorter than 10 kb (“RS3_short”). (E) Replication fork direction at the lower and upper breakpoints of SVs assigned to rearrangement signature RS1. Categories indicate the fork direction at each breakpoint (e.g., L–R = leftward at the lower breakpoint and rightward at the upper). Counts are normalized to expectations from randomly shuffled SVs matched for size. ^5^

We visualized “repli-seq”-derived replication timing for SV sets attributed to common non-clustered SV signatures, which included rescaling coordinates centered at SV midpoints (Figure 2C; Figure S2 shows unscaled analysis). Among these, SVs attributed to signature RS1—dominated by non-clustered tandem duplications in the 100 kb–1Mb size range—occurred in the earliest-replicating regions of the genome. Additionally, replication timing peaked at midpoints of SVs attributed to this signature, consistent with replication origin enrichment. For RS1, replication timing was significantly earlier at SV midpoints compared to SV margins (P < 1 *×* 10^-15^, Wilcoxon test), where margins were defined as regions within 1 Mb on either side of each SV. At finer resolution, replication timing at RS1 midpoints was also earlier than at SV breakpoints (P = 0.02, Wilcoxon test). Longer tandem duplications attributed to the RS14 signature—with a median size of 699 kb—were often found in a similar configuration, with midpoints replicating earlier than their margins (P < 1 *×* 10^-15^, Wilcoxon test). In contrast, randomly simulated SVs preserving the observed length distribution showed no enrichment of earlier replication timing at SV midpoints (Figure S2).

Earlier replication timing at SV midpoints indicates that replication origins are located within SVs rather than at the breakpoints. Indeed, a two-sided proportion test showed a significant enrichment of SNS-seq–defined replication origins^25^ within RS1 tandem duplications compared with flanking regions (rate ratio = 1.24; P < 1 *×* 10^-15^). In visualizations, the replication origins most commonly coincided with SV midpoints (Figure 2D). RS3, which includes tandem duplications in the 1-10kb and 10-100kb channels, shows a similar level of enrichment (rate ratio 1.32, P < 1 *×* 10^-15^), but the signal was specific to the 10-100kb duplications and disappeared for the 1-10kb duplications (Figure S3). In contrast, RS8—dominated by foldback inversions smaller than 10 kb—showed the opposite trend: fewer replication origins within the SV footprint compared to margins (rate ratio = 0.6, P < 1 *×* 10^-9^), suggesting that origin-poor regions are prone to inversion formation.

We further assessed the direction of the replication fork at both breakpoints of each SV, as a change in replication fork direction between breakpoints indicates the presence of a replication origin or termination zone within the spanned segment. We quantified fork direction using the dominant replication direction inferred from “OK-seq”,^28^ an assay applied to the HeLa cervical-cancer-derived cell line. Of note, these “OK-seq” data were independent of the replication timing and origin datasets used above. Tandem duplications attributed to RS1 showed a significant excess of L–R configurations (Figure 2E), characterized by leftward replication at lower breakpoints and rightward replication at upper breakpoints (OR = 1.08, P = 0.04). Compared to the abundance of randomly simulated L-R regions of matching sizes in the genome, the ratio of observed L-R duplications to expected reached 1.5 (see’Normalization of RFD Signatures’ in Methods). This effect was consistent across RS1 SVs in cancers from six tissue types where this signature is abundant (Figure S5).

For RS1 (100 kb–1 Mb duplications), we observed the enrichment of replication origins, earlier replication timing between SV breakpoints, and the predominant L–R replication fork direction at breakpoint loci. Together, these features are consistent with the SFBF mechanism for RS1-type tandem duplications (Figure 1A).

### APOBEC strand bias at SV loci supports bi-directional replication

Replication timing and fork direction are inherently stochastic,^25^ and replication profiles inferred from cell lines represent average behavior. An ideal tracer would instead capture replication fork orientation at the moment of SV formation. We posit that strands of adjacent APOBEC-attributed point mutations provide such a tracer. First, APOBEC mutations arise in a distinctive TCW context, which is readily identifiable.^29^ Second, their reported reference allele (C>N or G>N) reflects whether cytidine deamination occurred on the reference or complementary DNA strand, respectively. Third, this strand assignment is systematically coupled to replication direction due to single-stranded DNA exposed on the lagging strand: in leftward-replicating regions, APOBEC mutations are predominantly reported as G>N, whereas in rightward-replicating regions they are predominantly reported as C>N.^27^ Together, these properties imply that APOBEC-induced mutations encode local replication fork direction.

If SVs arise from stalled replication forks, they should be accompanied by predictable APOBEC strand patterns. Under the SFBF model, replication initiates bidirectionally from a single origin, and forks diverge. In this setting, APOBEC mutations are expected to accumulate interior to the SV breakpoints when mapped to the reference genome, with G-reported mutations near the lower (5′) breakpoint and C-reported mutations near the upper (3′) breakpoint (a “Gs-then-Cs” pattern; Figure 3A left). By contrast, SVs generated by converging forks should show APOBEC mutations accumulating exterior to the SV, with the opposite “Cs-then-Gs” orientation (Figure 3A right), providing a testable distinction between models. In both scenarios, prolonged exposure of single-stranded DNA at stalled forks is expected to result in an excess of APOBEC mutations proximal to SV breakpoints.

**Figure 3.**
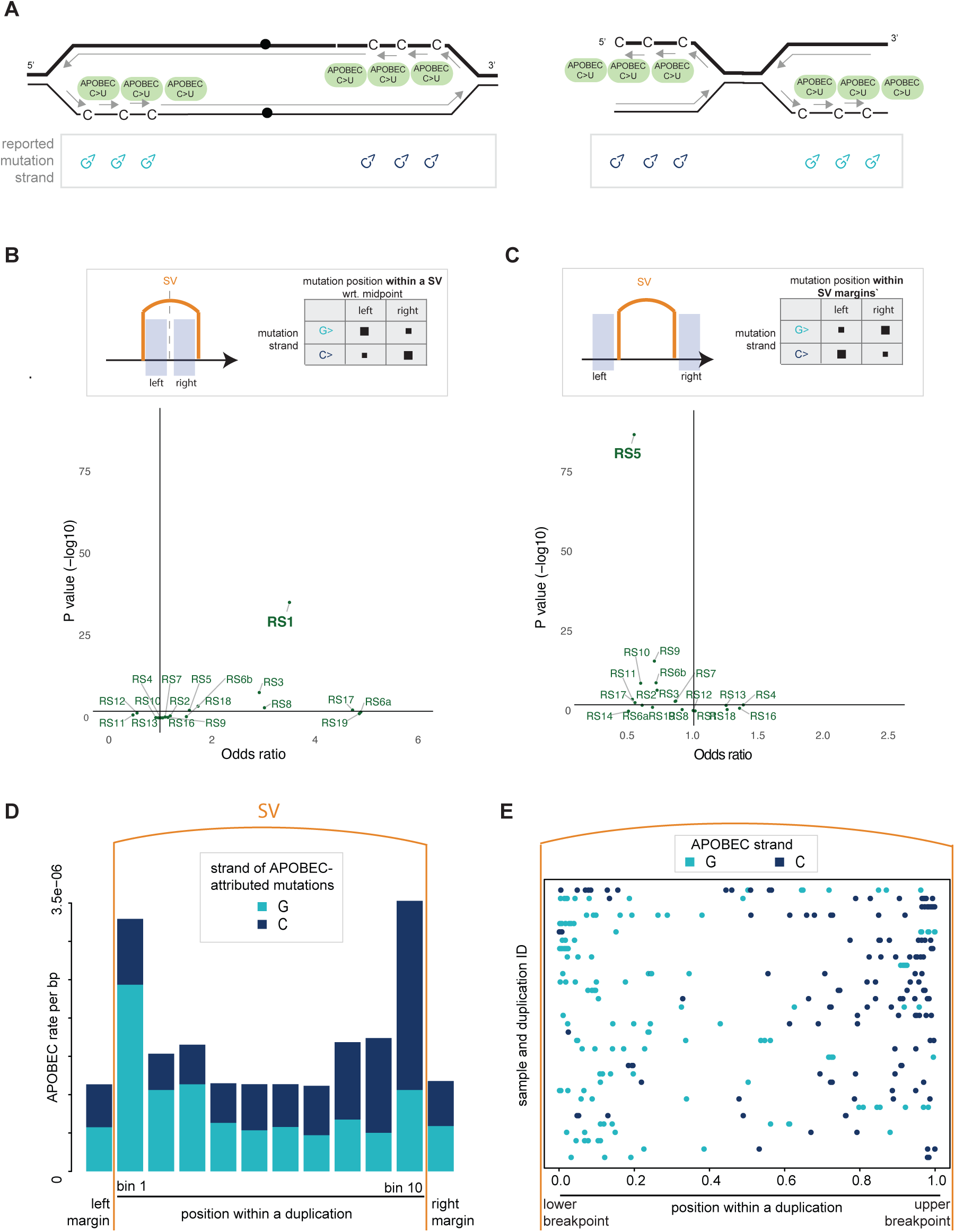
Replication fork direction inferred from APOBEC strand bias near structural variants (SVs). (A) Schematic illustrating the expected locations and strands of APOBEC-induced mutations near stalled replication forks. Left: two replication forks progressing bidirectionally from a single origin. Right: converging forks arriving from two distinct origins. (B) Association between the strand of APOBEC-attributed mutations overlapping SVs and their position along the SV span. RS1, a signature dominated by 100 kb–1 Mb tandem duplications, exhibits a pronounced strand bias, with G-reference mutations enriched near lower breakpoints and C-reference mutations near upper breakpoints (Fisher’s exact test; 2×2 contingency table). (C) Strand bias of APOBEC-attributed mutations located within 100 kb flanking, but non-overlapping, margins of SVs. RS5, a signature enriched for 1–10 kb deletions, shows the opposite pattern: C-reference mutations predominate in left margins and G-reference mutations in right margins. (D) Metaprofile of APOBEC-attributed mutation strand (C- or G-reference) across RS1 tandem duplications and ±100 kb flanking regions in PCAWG samples. (E) Representative examples of APOBEC-attributed mutation patterns w7ithin RS1 tandem duplications, limited to SVs containing at least five such point mutations.

To test these predictions in the PCAWG dataset, we quantified the strand bias of APOBEC-attributed mutations occurring within SVs and in their flanking regions. Mutations were extracted from SV-overlapping regions and from 100-kb windows immediately adjacent to each SV. To exclude APOBEC-attributed mutations that predated SV formation, we restricted the analysis to mutations with multiplicity of 1, and verified that relaxing this filter did not alter the conclusions (see Methods). A subset of SVs overlapped with APOBEC mutations: for example, 1,675 of 26,417 (6.3%) RS1-attributed events harbored at least one APOBEC mutation, accounting for 1,853 of 370,790 APOBEC mutations (0.5%) in this group of 193 samples (Figure S6). For the sets of SVs attributed to RS1 and the remaining signatures, we compared the proportion of reference G- versus C-reported mutations between the left and right halves, using odds ratios to quantify strand asymmetry, with OR>1 indicating the “Gs-then-Cs” pattern. The strongest bias was observed for RS1 tandem duplications (OR = 3.5, P < 1 *×* 10^-16^; Figure 3B; Fisher’s exact test), consistent with leftward fork polarity at the lower breakpoint and rightward at the upper. For mutations occurring in the regions immediately adjacent to SVs, the proportion of APOBEC mutations in RS1 margins displayed no significant strand preference (OR = 1.02; Figure 3C). In contrast, the deletion-rich signature RS5 displayed a reversed bias in SV margins (OR = 0.3, P < 1 *×* 10^-16^), with C-predominating on the left and G-reported mutations on the right. We thus observe opposing strand biases of APOBEC-attributed mutations around SVs of different types.

To resolve the spatial distribution of APOBEC activity within RS1-associated duplications, we analyzed the mutation density at higher resolution across SV bodies. We divided each SV into ten equal bins and computed per-base, per-bin APOBEC mutation rates across bins aggregated across SVs and samples (Figure 3D). APOBEC mutations were highly enriched near breakpoints within duplicated regions, with rates over threefold higher than at SV midpoints (P < 1 *×* 10^-15^, Poisson test), consistent with elevated APOBEC activity at replication fork stalling sites. Moreover, APOBEC mutation density at breakpoints exceeded that in regions flanking the SVs, indicating a localized burst of APOBEC activity coincident with SV formation rather than background mutagenesis before or after the event. Strand bias was also breakpoint-localized: 74% of mutations were G-reported near lower breakpoints and 30% near upper breakpoints (Poisson 95% CIs: 64–84% and 24–37%, respectively), recapitulating the “Gs-then-Cs” pattern described above.

Inspection of individual RS1 events confirmed this aggregate trend. In SVs with at least five APOBEC-attributed mutations, APOBEC mutations were concentrated near both breakpoints, with G-reported mutations predominating near the lower breakpoints and C-reported mutations near the upper breakpoints (Figures 3E, S7). Some exceptions were observed in duplications embedded within compound SVs, which can obscure strand assignment (Figure S8). Notably, APOBEC mutations were spaced >1 kb apart, suggesting a mechanism distinct from kataegis.^24^ Altogether, the APOBEC patterns in individual duplications indicate that the formation of larger tandem duplications involved bidirectional replication, with forks diverging from an origin placed near the center of the TD segment. These patterns support the SFBF model.

A comparable APOBEC strand pattern was observed in *templated insertions*. Templated insertions are semi-reciprocal interchromosomal translocations that generate copy-number gains.^6,30^ In these events, G-reported mutations appeared near lower breakpoints, and C-reported mutations near upper breakpoints, mirroring the strand configuration of tandem duplications and supporting a shared replication-origin mechanism (Figure S9). The shared patterns of APOBEC strand observed for simple TDs and templated insertions suggest that these two types of SVs arose from a common SFBF mechanism. A common replication–origin–driven mechanism appears to generate localized segmental gains through both simple and compound SVs.

### Support for the SFBF model in *CCNE1*-amp and *CDK12-/-* tumors

We next examined whether bi-directional replication at TDs and the SFBF model are supported specifically in tumors with *CCNE1* amplification or bi-allelic *CDK12* loss. This analysis was motivated by prior reports suggesting HRD in these therapeutically challenging populations.^31–34^ Although tumors with these cell-cycle alterations in the PCAWG cohort exhibited the HRD-associated point mutation signature SBS3, they lacked the composite HRD phenotype, including its characteristic indel and SV components (Figure S10; Supplementary Note). Instead, *CCNE1*-amplified tumors were enriched for 100 kb–1 Mb duplications (RS1), whereas *CDK12*-deficient tumors displayed similar and longer non-clustered TDs (RS1, RS14; Figure 4A). Accordingly, TD length distributions across *BRCA1*-mutant, *CCNE1*-amplified, and *CDK12*-deficient tumors were tri-modal, with alteration-specific peaks (Figure 4B),^5,9,10^ supporting HRD-independent mechanisms of TD formation.

**Figure 4.**
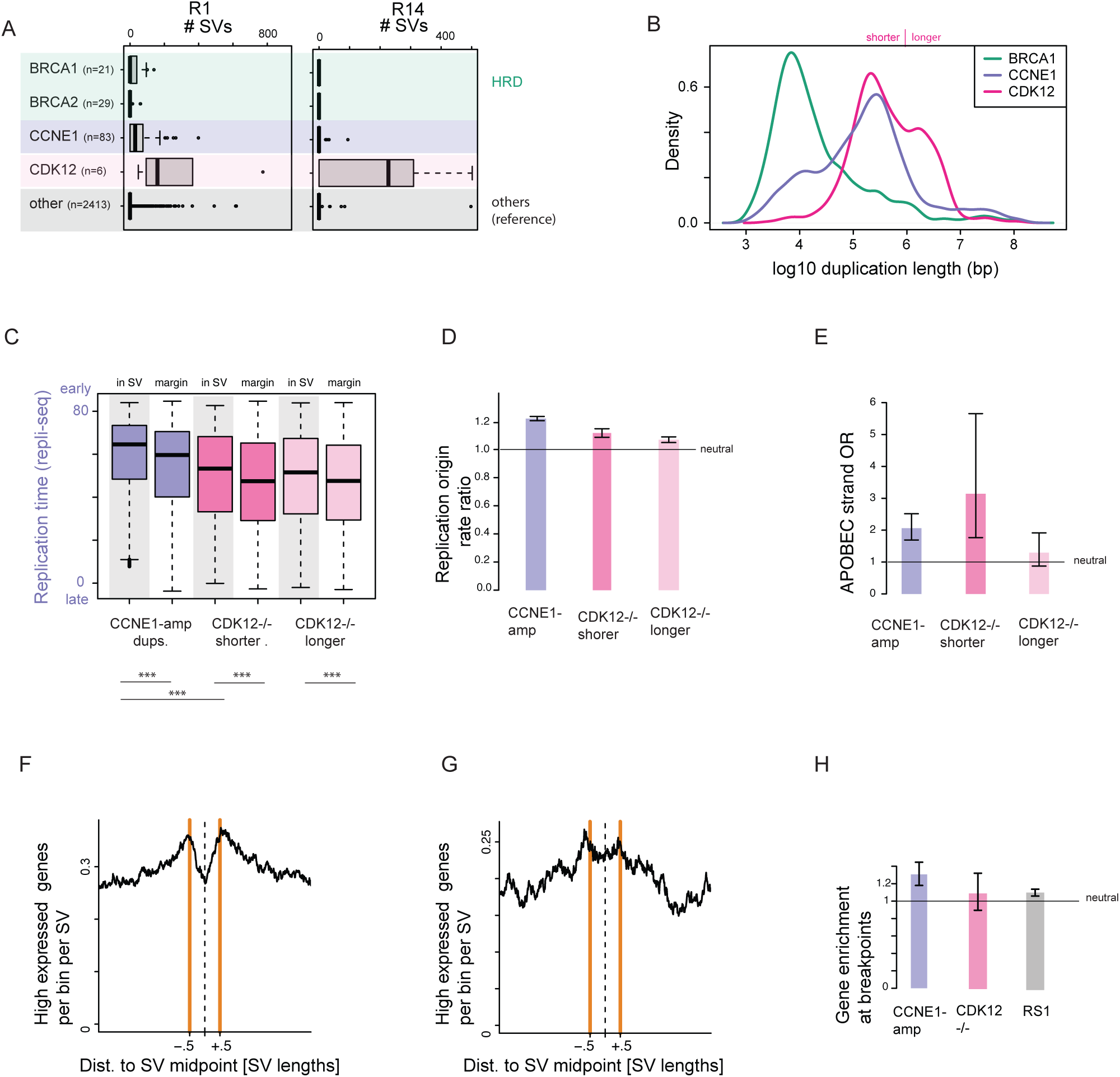
Genomic features of tandem-duplication loci in CCNE1-amplified and CDK12-mutant cancers. (A) *CDK12*-mutant tumors are enriched for medium-and long-range tandem duplications (RS1, RS14), whereas *CCNE1*-amplified tumors predominantly show RS1-type rearrangements. (B) Tandem-duplication lengths in *BRCA1*-deficient, *CCNE1*-amplified, and *CDK12*-mutant cancers display a trimodal distribution. (C) Replication time (repli-seq) at tandem-duplication loci, stratified by gene variant and compared to proximal regions (margins). (D) Enrichment of replication origins (SNS-seq) overlapping tandem duplications compared to SV margins, stratified by gene variant. 95% CIs from a proportion test. (E) Correlation of APOBEC strand with leftward and rightward orientation with respect to the tandem-duplication midpoint, stratified by gene variant. Tandem duplications in *CDK12*^-/–^ tumors are further subdivided by whether they exceed 1 Mb in size. Odds ratios and 95% CIs from Fisher’s exact test. (F) In in *CCNE1*-amplified tumor, highly expressed genes coincide more strongly with tandem-duplication breakpoints than with midpoints. (G) Profiles of highly expressed genes around tandem duplications in *CDK12*^-/-^ tumors. (H) Quantification of the coincidence of highly expressed genes with SV breakpoints compared to the midpoint. Fisher’s exact test and 95% CIs; random SVs matched for length used as controls.

To probe SV mechanisms in *CCNE1*-amplified tumors, we analyzed replication origins, timing, direction, and transcription around duplication loci. We observed an enrichment of replication origins (rate ratio 1.22, P < 1 *×* 10^-16^, proportion test comparing SV segments flanking regions; Figure 4D) and earlier replication timing at duplication midpoints relative to breakpoints (P = 0.002, Wilcoxon test; Figure 4C). APOBEC-attributed point mutations followed the expected strand pattern, with G-reported mutations enriched 3’ of lower SV breakpoints and C-reported mutations enriched 5’ of upper SV breakpoints (Figure 4E; OR = 2.1, P < 1 *×* 10^-12^, Fisher’s exact test; considering RS1 attributed TDs; Figure 4E). Overall, larger duplications in *CCNE1*-amplified cancers appear centered on replication origins, consistent with bidirectional forks diverging from a central origin.

In *CDK12*-deficient tumors, tandem duplications also showed earlier replication timing at SV midpoints (P = 0.03) compared to the breakpoints and were enriched for replication origins near midpoints (rate ratio 1.19, P < 1 *×* 10^-16^; Figure 4C). APOBEC strand bias in shorter (<1 Mb) *CDK12* duplications matched the RS1 pattern (OR = 3.1, P < 1 *×* 10^-4^, Fisher’s exact test; Figure 4E), but was weaker for longer duplications (OR = 1.3, P = 0.18), likely reflecting signal dilution due to the larger genomic footprint and background APOBEC activity. While consistent with the SFBF model, tandem duplications in *CDK12*-deficient tumors occur in later-replicating regions than those in *CCNE1*-amplified tumors (median replication timing 60.3 vs 51.4; P < 10^-15^ in Wilcoxon test; Figure 4D).

We next examined replication-transcription collisions as a possible cause of fork stalling. Most SVs overlapped with protein-coding genes: 76% of tandem duplications in *CCNE1*-amplified tumors and 68% in *CDK12*–/– tumors had at least one breakpoint coinciding with a gene (see Methods). Across all RS1 duplications, SV breakpoints were more likely to intersect highly expressed genes than midpoints (OR = 1.12, P = 1 *×* 10^-9^; Fisher’s exact test, genes expressed above median in ovarian cancer, when compared to randomly matched SVs with same size distribution), a pattern not observed for lower expressed genes (Figure S2). This enrichment was particularly pronounced for *CCNE1*-amplified tumors (OR = 1.28; P = 1 *×* 10^-6^; Figures 4F, 4H) and more modest in *CDK12*-deficient tumors (OR = 1.09; not significant; Figures 4G, 4H). In *CCNE1*-amplified tumors, tandem duplication breakpoints additionally showed a gene-strand bias: lower breakpoints overlapping genes tended to align with the positive strand, and upper breakpoints with the negative strand (OR = 0.69, P < 1 *×* 10^-4^, Fisher’s exact test; Figure S11). This strand configuration suggests head-on transcription–replication encounters. For tandem duplications in *CDK12*-/- tumors, the strand association was near neutral (OR=1.01) and not significant. Together, these correlations implicate replication–transcription collisions at tandem duplication loci in tandem duplication formation, whose S phase timing differs between *CCNE1*-amplified and *CDK12*-mutant cells.

### Expanding SV signatures with replication features

Re-analysis of PCAWG SVs assigned to known mutational signatures revealed striking biases in replication direction and timing—features not yet integrated into algorithms for SV signature extraction. To better characterize known signatures and identify new ones, we first incorporated replication-fork direction (RFD) as an additional feature, extending the standard 16 SV categories^24^ (defined by SV type and inter-breakpoint distance). Each SV was further classified according to local RFD at its lower and upper breakpoints (L–L, L–R, R–L, R–R), yielding 64 replication-direction–aware features.^24^

From 577 PCAWG samples spanning breast, ovarian, prostate, lung, and esophageal cancers—each with at least 30 non-clustered SVs—we extracted RFD-based signatures. Using MuSiCal’s stability criteria,^35^ we identified 15 stable signatures, six of which were dominated by non-clustered tandem duplications (Figures 5A, S12). For visualization, we normalized signatures to correct for the genomic opportunity of each replication-fork direction configuration (see Methods). As expected, duplication-dominated signatures in the 100 kb–1 Mb (RFD9) and 1–10 Mb (RFD8) ranges were enriched for the L–R replication channel (leftward at the lower breakpoint, rightward at the upper).

**Figure 5.**
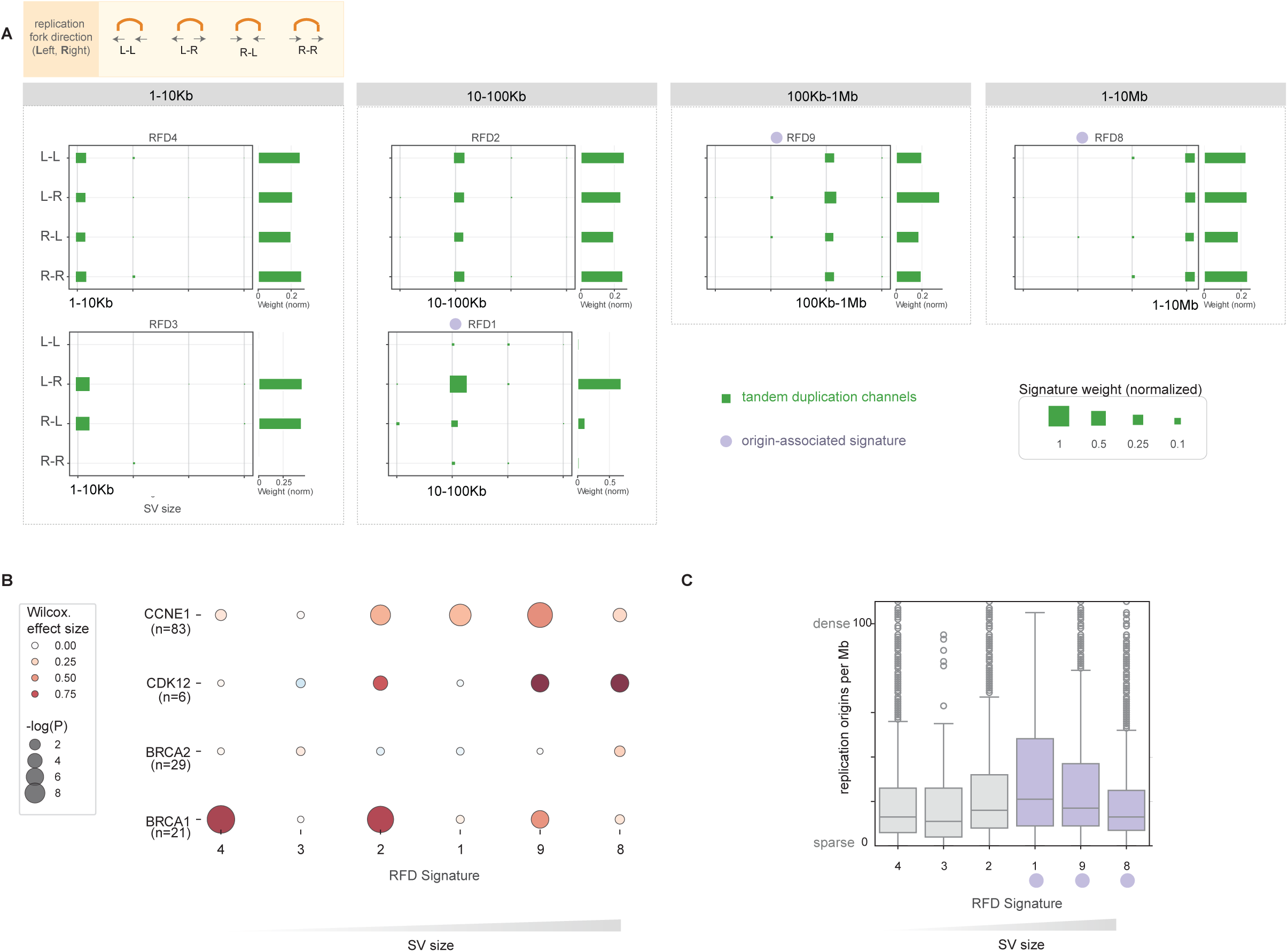
Expanded structural variant (SV) signature discovery in the PCAWG dataset. (A) De novo discovery of SV signatures incorporating replication-fork direction (RFD) at breakpoints reveals new splits among tandem-duplication signatures. Tandem duplications of 1–10 kb dominate signatures RFD3 and RFD4, whereas those of 10–100 kb dominate RFD1 and RFD2. Barplots on the right show row sums of shown non-clustered TD columns. Complete signatures are visualized in Figure S12. (B) Enrichment of RFD signatures across tumor groups defined by *CCNE1* amplification or loss of *CDK12*, *BRCA1*, or *BRCA2*. Effect sizes and P values were computed using the Wilcoxon rank-sum test. (C) SVs were assigned to their most likely RFD signature. Comparing SVs assigned to RFD1 (10–100 kb), RFD9 (100 kb–1 Mb), and RFD8 (1–10 Mb), increased SV size is associated with decreased density of replication origins.

Unexpectedly, we identified two distinct duplication signatures in the 10–100 Kb range: RFD1 was specifically associated with L-R loci, while RFD2 displayed no such association. Furthermore, while both RFD4 and RFD3 were dominated by the shortest tandem duplications within the 1-10Kb range, they differed by replication direction biases: RFD4 duplications were enriched for events with consistent direction across breakpoints (L-L, R-R), while RFD3 duplications often occurred at loci where the direction switched (L-R, R-L). Together, these splits suggest that shorter tandem duplications may arise from at least two distinct mechanisms, one potentially linked to origin-associated processes like SFBF.

RFD-defined signatures showed distinct associations with alterations in genes involved in DNA repair and replication stress. Signature exposures were compared across *BRCA1*-/-, *BRCA2*-/-, *CDK12*-/- and *CCNE1*-amplified tumors relative to the remaining samples. For example, both RFD1 and RFD2 signatures comprised mainly non-clustered 10–100Kb tandem duplications but differed in their associations: RFD1, enriched for L-R events, was specifically linked to *CCNE1* amplification (Figure 5B), whereas RFD2 was additionally associated with loss of *CDK12* and *BRCA1*.

Across three SV size ranges, distinct replication fork directionality (RFD) signatures were enriched at L–R loci: RFD2 (10–100 kb), RFD9 (100 kb–1 Mb), and RFD8 (1–10 Mb), all putatively associated with replication origins. Based on sample-level exposures and features of individual SVs, we assigned SVs to these newly defined RFD-based signatures. Each SV was further characterized by local replication origin density. Across the SV groups attributed to L-R associated signatures RFD1, RFD9, and RFD8, increasing SV size was associated with reduced local replication origin density (Figure 5C). Assuming that these SVs arise at replication origins, the observations are consistent with a model in which tandem duplication length for these larger TD types is determined by the distance traveled by replication forks diverging from a single origin. The size of larger TDs may thus be a reflection of replicon size.

Motivated by the observation that TD loci in *CDK12-/-* tumors are typically replicated later than those in *CCNE1-amplified* tumors, we next asked whether replication timing at SV midpoints would reveal additional signature structure. To address this, we performed an independent signature discovery experiment using replication timing (RT) features. Each non-clustered SV was subclassified by repli-seq timing at its midpoint (very early, early, mid, late, very late), expanding the 16 SV categories to 80 timing-aware features. This analysis identified 18 RT signatures with an average silhouette score above 0.7 (Figure S12). We discovered multiple signature splits along the new RT dimension. For example, 100Kb-1Mb duplications split into two signatures: one dominated by SV at earlier and one at later replicating loci (RT1 and RT2, respectively). Similar splits were observed for 1-10Kb inversions (RT9 and RT10) and translocations (RT7 and RT8). Taken together, these results demonstrate that replication timing and direction shape the SV mechanisms and population-level patterns.

### SVIG: a multi-class classifier of DNA repair deficiencies and replication stress

Signature splitting upon incorporation of replication fork direction (RFD) and replication timing (RT) indicates that these features capture previously underappreciated population heterogeneity. We next examined the population structure defined by these refined SV features. To assess their utility for predicting repair and replication stress phenotypes, we used RFD- and RT-informed signature exposures from PCAWG breast, ovarian, prostate, esophageal, and lung cancers to train a multi-class classifier (Figure 6A). The classifier was termed SVIG for “Structural Variant-based Inference of Genotype”. Samples were labeled according to previously defined DNA repair and cell cycle genotypes (*BRCA1*-/-, *BRCA2*-/-, *CDK12*-/-, and *CCNE1*-amp), and SVIG was trained to predict these classes from SV signature exposures. SVIG is based on a multi-class XGBoost algorithm with class-weighted training to correct for sample imbalance (see Methods). Differential weighting of sample classes mitigates class imbalance and label imperfection arising from undetected molecular alterations. Newly derived replication fork direction and timing features are among the most important SV features learned by the classifier (Figure S13).

**Figure 6.**
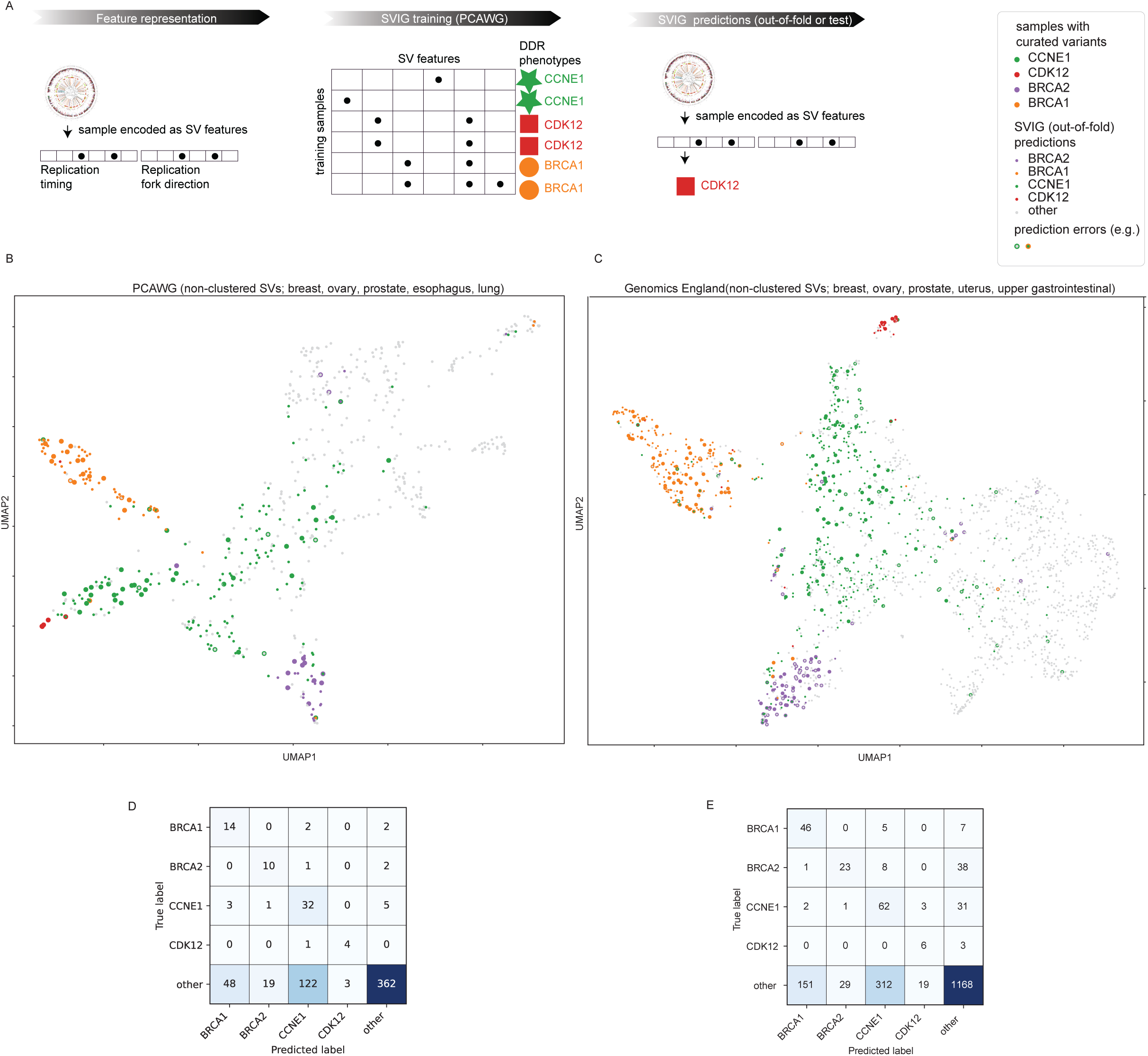
Unsupervised and supervised analysis of expanded structural variant (SV) signature exposures. (A) Diagram showing representation of cancer samples as SV features, SVIG training, and application to test and out-of-fold samples (B) PCAWG cohort. Left: UMAP of SV-signature exposures highlighting tumors with loss-of-function mutations in *CDK12*, *BRCA1*, or *BRCA2*, and those with *CCNE1* amplification (large dots). Smaller dots represent out-of-fold predictions from a multi-class XGBoost classifier trained to distinguish these alteration groups. (C) Genomics England cohort. This independent dataset includes a larger number of samples with the alterations of interest. The PCAWG-trained classifier was applied directly, without retraining or additional tuning. (D) Confusion matrix showing out-of-fold classifier performance in the PCAWG dataset. (E) Confusion matrix for classifier performance in the Genomics England dataset, unseen during training.

We visualized the input signature exposure data and out-of-fold predictions on the PCAWG dataset (generated on samples held out during training) using UMAP projections and confusion matrices (Figures 6B, 6D). UMAP was applied to normalized signature exposure vectors. The classifier showed strong recall for all four classes (*BRCA1*-/-: 0.78, *BRCA2*-/- 0.77, *CCNE1*-amp: 0.78, *CDK12*-/-: 0.8); precision was lower (*BRCA1*-/-: 0.22, *BRCA2*-/-: 0.33, *CCNE1*-amp: 0.20, *CDK12*-/- 0.57). The lower specificity, when assessed against variant-defined labels, suggests that these SV phenotypes are not restricted to tumors with known alterations, implying additional mechanisms of pathway inactivation. Among the tumors with predicted’*CCNE1*-amp-like’ phenotype but without a conservatively defined *CCNE1* amplification, we observed an enrichment of *TP53* mutations (OR = 6.4, P < 1 *×* 10^-14^), amplifications of *YWHAZ* gene in proximity of the *MYC* locus (OR = 6.7, P < 1 *×* 10^-4^), *MYB* amplifications (OR not estimable, P < 1 *×* 10^-4^), and loss-of function variants in *CDKN2A* (OR = 5.0, P < 1 *×* 10^-4^) and *CDKN2B* (OR = 3.0, P < 1 *×* 10^-4^, two-sided Fisher’s exact test). Among the samples classified as’*CCNE1-like*’, there was no apparent separation on the UMAP between samples carrying *CCNE1* amplifications (Figure 6B). SVIG identifies SV phenotypes consistent with replication stress, some of which may arise via alternative genomic mechanisms beyond *CCNE1* amplifications.

We next applied the pre-trained SVIG to the Genomics England (GEL) dataset,^36^ an independent and much larger cohort (n=1915) of breast, ovarian, prostate, uterine, and upper gastrointestinal tumors. UMAP visualization of SV exposures and predictions (Figures 6C, 6E) showed that recall was comparable to that achieved for out-of-fold samples in the PCAWG set: for *BRCA1*-/- (0.79), *CCNE1*-amp (0.63) and *CDK12*-/- (0.67). Recall for the *BRCA2*-/- group was markedly reduced (0.33). In particular, 38 out of 70 samples from the GEL dataset with confirmed *BRCA2* loss were misclassified as “other.” Retraining SVIG on the GEL dataset improved out-of-fold accuracy across all classes, including the *BRCA2*-/- (recall 0.87) and *CCNE1-amp*-amp (recall 0.81) classes, highlighting additional predictive power gained from the larger dataset. SVIG provides molecular interpretation of SV landscapes, confirming or revealing DNA repair and replication stress phenotypes, even if the responsible genetic variants are not detected.

## DISCUSSION

To better understand and quantify replication stress, we described the distinct relationships between SVs and DNA replication timing, direction, and origins. In particular, larger tandem duplications frequently coincide with replication origins, which are enriched at SV midpoints rather than breakpoints. This pattern is consistent with the SFBF model, in which a TD arises from bidirectionally collapsing forks emerging and diverging from an origin. The enrichment of larger (> 100 kb) duplications near replication origins and their APOBEC strand asymmetry both argue against aberrant fork restart or MMBIR as the primary sources of longer SVs, because neither of these mechanisms entails bidirectional replication from origins (Figure 7).

**Figure 7.**
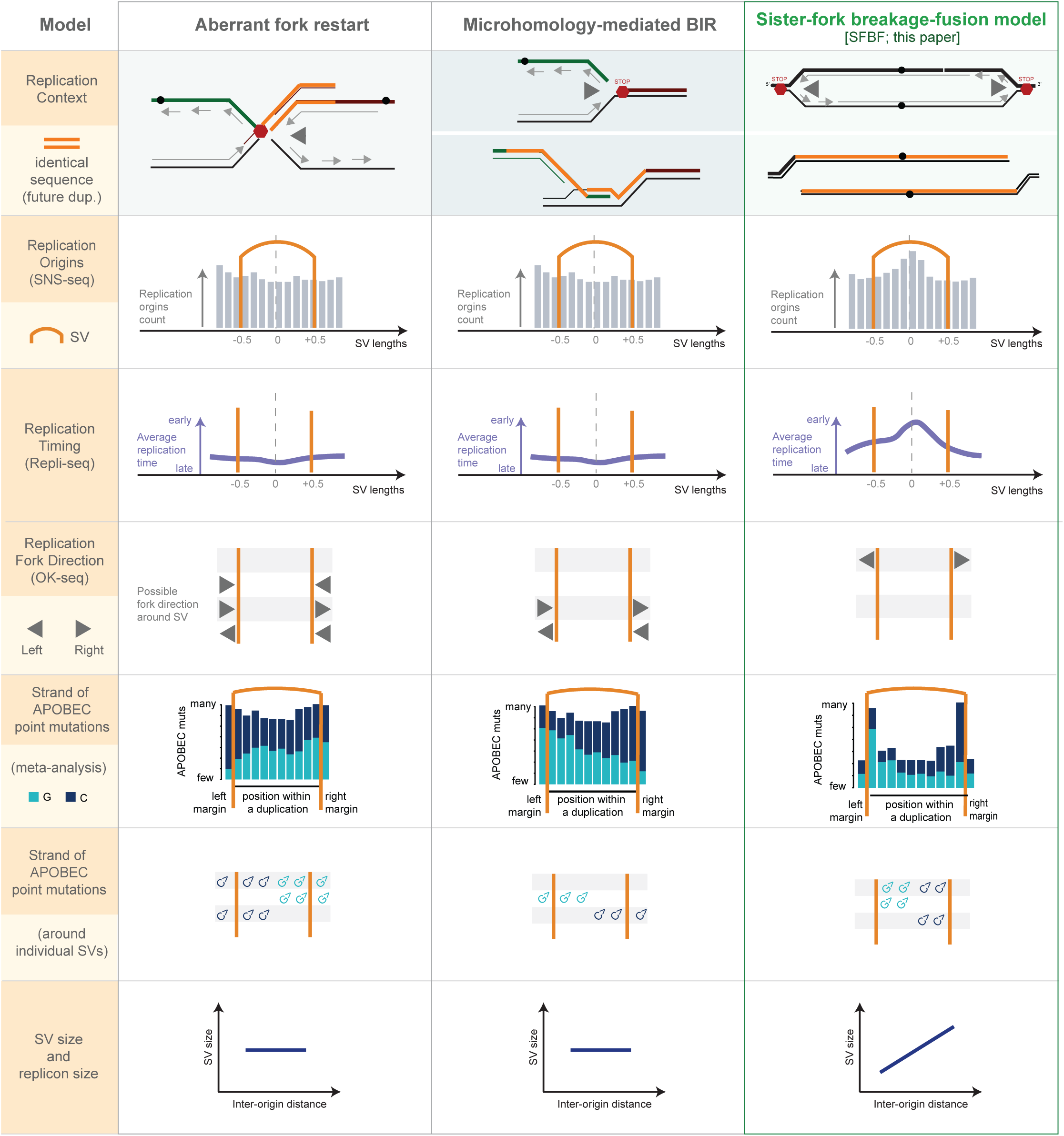
**Mechanistic models for tandem duplication formation and their predictions for replication origins, fork directionality, and APOBEC mutation strand bias examined in this study.**

Our data support at least two mechanisms for tandem duplications generating SVs with distinct size distributions. Small (<10 kb) tandem duplications in *BRCA1^-/-^*tumors have been linked to aberrant fork restart^19^ or microhomology-mediated break-induced replication.^20,37^ In contrast to larger (> 100 kb) TDs where our data supports the SFBF model, smaller duplications do not show origin enrichment (Figure S3) and their APOBEC strand patterns are inconclusive. Our signature analysis further suggests that there is no absolutely defined boundary between these distinct TD mechanisms; the zone of TD size between 10 and 100 kb is a region where both BRCA1-linked TD mechanisms and SFBF mechanisms may operate.

Further details of the SFBF model remain to be elucidated. We propose that it involves an end joining step, which requires experimental validation. Of note, the short microhomologies observed at tandem duplications (average 1.8 bp for RS3 TDs, 1.9 bp for RS1, and 2.1 for RS14; Figure S4) are inconsistent with classical MMBIR, which requires longer tracts of identical sequences for strand invasion.^3^

It remains unclear why cancers with *CDK12* loss and *CCNE1* amplification exhibit large tandem duplications centered at replication origins. One hypothesis relates to impaired replication origin licensing, as demonstrated in *CCNE1*-amplified tumors^38^ and possibly extending to *CDK12*-deficient tumors.^39^ Prior studies suggest that *CDK12* and Cyclin K are required for efficient loading of pre-replication complexes onto chromatin.^39,40^ In cells with defective origin licensing, replication forks traverse longer distances, increasing their vulnerability to stalling. Additional mechanisms may contribute in *CDK12*-/- tumors, including transcription–replication conflicts and R-loop DNA–RNA hybrids,^41–43^ reflecting *CDK12*’s roles in transcriptional elongation, termination, 3′ end processing, and mRNA splicing.^44^ Indeed, TDs in both *CDK12*-mutated and *CCNE1*-amplified tumors frequently localize to highly transcribed regions, but the replication–transcription collision signatures are more pronounced in *CCNE1*-amplified tumors. Independently of transcription, the excess tandem duplications in *CDK12*^-/-^ cells could result from aberrant origin relicensing and origin re-firing,^21^ as part of the SFBF mechanism. However, this hypothesis is untested in human cells.^3^

Other SV classes likewise align with the features of DNA replication. Gains from semi-balanced translocations (“templated insertions”^6^ or “synthetic duplications”^30^) share APOBEC strand patterns with tandem duplications (Figure S9), consistent with stalled bidirectional forks that resolve in a manner alternative to tandem duplications. Their size correlation with tandem duplications, as observed previously,^30^ supports a shared replication origin–based mechanism. On the contrary, short deletions (<10 kb, RS5) display opposite APOBEC strand biases near deleted regions, suggesting incomplete replication by converging forks.

Despite using three complementary approaches to quantify replication dynamics, our analysis is constrained by the inherent stochasticity of human DNA replication and the limited number of SVs per tumor. In the absence of patient-matched replication profiles, we relied on Repli-seq data from the MCF7 cell line, which has been shown to resemble other ENCODE cell lines, with 60% of the earliest-replicating regions shared across lines.^27^ Consistent with this, we replicated the association between larger TDs and early replication timing at SV midpoint, replication origins, and bidirectional replication across cancer types (Figure S5). Although reproducible, replication-timing and fork-direction signals around SVs exhibit only moderate effect sizes. In contrast, APOBEC strand asymmetry provides a more direct readout of replication direction—likely at the time of SV formation—and yields stronger effect sizes. Future studies may leverage engineered cell lines with abundant SVs and elevated APOBEC activity to address the current limitation that only a minority of SVs harbor APOBEC mutations. As whole-genome sequencing datasets expand, particularly those enriched for tumors with DNA-repair and cell-cycle defects, SV signature analyses will gain statistical power.

Our findings suggest that SVs capture a signature of replication stress and may help identify patients likely to benefit from drugs that exploit replication stress as a vulnerability of cancer cells.

## Supplemental Tables

1. Table 1: Replication, transcription, and repeat features of SV signatures as defined in Degasperi et al.
2. Table 2: APOBEC strand polarity, within and in margins of SVs, for SV signatures as defined in Degasperi et al.
3. Table 3: Redefined SV signatures with replication fork direction (RFD) and timing (RT) features, and exposures in the PCAWG set.
4. Table 4: Sample labels and SVIG predictions on the PCAWG dataset.

## Supplementary Note

### Assessment of HRD signatures of *CCNE1*-amplified and *CDK12*-mutated cancers

While the signature of homologous recombination (HR) deficiency (HRD) has been described in tumors with *BRCA1/2* loss, there is conflicting evidence whether alterations in cell cycle-related genes (*CCNE1*, *CDK12*) produce comparable HRD-associated signatures.^31–34^ HRD is marked by characteristic mutational signatures across variant types: SBS3 (SNVs), ID6/ID8 (indels), and RS3/RS5 (SVs) (Figure S10).^2^ We verified *CDK12*-deficient tumors from the PCAWG cohort lack the composite HRD signature in its entirety.

They sometimes harbor SBS3 point mutations but generally lack HRD-associated indels and SVs (n=6 PCAWG samples; 1 sample with both *CDK12* loss and *CCNE1* amplification not considered). Consistently, HRDetect predicted HR proficiency in all seven *CDK12* bi-allelic loss-of-function tumors (probabilities of HR deficiency 0.00–0.07, well below the 0.70 threshold).^2^ Similarly, *CCNE1*-amplified tumors lack the indel and SV components of the HRD signature (Figure S10; n=86 PCAWG samples). Of 83 *CCNE1*-amplified cancers with HRDetect data, 71 were classified as HR proficient.

## METHODS

### Datasets

#### Pan-Cancer Analysis of Whole Genomes (PCAWG) Dataset

We accessed somatic SV calls from the collection of 2,778 cancer genomes across 39 types. We used the consortium’s consensus SV calls dated’161116’. The PCAWG consensus callset consists of SVs called by at least 2 of the following 4 callers: SvABA, DELLY, BRASS and dRanger.^6^ In the analysis, we used data from 1,797 samples with consensus SV calls and SV signature exposures (see below). In total, we analyzed 246,694 SVs. For APOBEC analysis, we used the consensus calls obtained from multiple single nucleotide variant callers produced by the PCAWG consortium, where evidence from at least 2 out of 4 callers (CaVEMan v1.5.1, MuTect, MuSE v1.0rc, DKFZ) was required per variant.^23^

#### Genomics England Cancer Dataset

We extracted non-clustered SV catalogs from GEL’s breast, ovarian, prostate, uterine, and upper gastrointestinal cancers. We used highly curated calls obtained by merging results from multiple callers, as published previously.^45^ After removing samples with insufficient numbers of SVs, the replication timing (RT) matrix included 4948 samples and 161 SV features, including a total of 430,517 SVs. The RFD matrix included 4948 samples and 129 features, and a total of 422,710 SVs.

#### Assignment of PCAWG SVs to signatures

We retrieved pan-cancer structural variant (SV) signatures (e.g. RS1-RS20) and their sample-specific exposures (i.e., signature abundances) from a previously published resource.^5^ We then attributed SVs to signatures by using the probability that a signature can generate a given type of SV, and the probability of a signature in a sample, corresponding to normalized exposures.

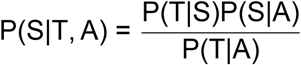

where:

- P(S|T, A) is the probability that the SV of type T was generated by mutational signature S in sample A.
- P(T|S) is the probability that a mutational signature S generated an SV of type T. This corresponds to known signature profiles, after normalization so that

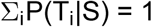

- P(S|A): probability of a signature generating a mutation in a given sample. Equivalent to normalized exposure of a signature in a given sample.
- P(T|A): probability of observing a SV of a given type in given sample. Obtained by marginalizing P(T|S)P(S|A).

For each SV, we attributed it to a signature that had the highest posterior probability of assignment (maximum-a-posteriori).

#### Assignment of PCAWG single nucleotide variants to signatures

We followed an equivalent procedure to assign point mutations to established mutational signatures. We used signatures and exposures calculated previously.^46^ We assigned each mutation to the signature with the highest posterior probability. For the APOBEC analysis, we used mutations attributed to signatures SBS2 and SBS13.

#### Analysis of SV positions with respect to replication origins, timing, and direction

The analysis of replication timing and replication origins within and around SVs was performed in two ways: (1) the SV region is rescaled by the length of each SV such that the SV has a unit length of 1 and we include 3 SV lengths on either side as a margin; (2) 1Mb around the SV midpoint on either side, irrespective of SV length. Rescaling in (1) allows us to easily relate replication origins to breakpoints of many SVs, each of which might be of different length length. (2) allows for a more direct comparison of region characteristics between SV groups of different sizes.

We utilized replication timing data for the MCF7 cell line from the ENCODE project,^26,27^ specifically the Wavelet-smoothed Signal, which represents a weighted average of newly replicated DNA across different phases of the cell cycle, followed by smoothing. Higher values of this metric correspond to earlier replication of a region, and lower values correspond to replication later in the cell cycle.^47^

For scenarios (1) and (2) we found regions with measured replication timing that overlapped with regions with SVs. For scenario (1) we worked with 100 bins per SV length, and for scenario (2), we worked with 100 bins per Mb. Within each bin around the SVs, we calculated the average replication timing, which we report in the summary curves in Figure 2C and Figure S2.

To characterize the replication origin landscape near structural variants (SVs), we used core replication origins identified by SNS-seq.^25^ This tissue-agnostic set accounts for approximately 80% of replication initiation events across 18 diverse human cell types.^25^ Using LiftOver tool,^48^ we remapped the origin positions to genome build hg19, and kept the core origins defined as those with origin firing frequency in two top deciles.

We found core origins overlapping with the genomic regions where SVs occurred. For each replication origin overlapping with an SV span or margin as defined above, we calculated its relative position to the SV midpoint. For all such SV-origin pairs, we calculated a histogram of relative positions of origins to SV midpoints. The histograms (gray barplots) are presented for two scenarios: (1) in Figure 2C, and (2) in Figure S2.

We described the dominant replication direction at SV breakpoints using data from the OK-seq assay. The OK-seq profiles Okazaki fragments characteristic of lagging replication and tags the strand of detected Okazaki fragments, such that the dominant replication direction in each region can be defined. We used the dominant replication direction measured in the HeLa MRL2 cell line.^28^ Each structural variant has two breakpoints when mapped to the reference genome. We overlapped breakpoint positions with dominant replication fork direction according to OK-seq profiles of HeLa cells. Replication fork direction around each SV was described as a tuple (D_1_, D_2_) where D_i_ *∈* {L, R}.

#### Analysis of SV positions with respect to transcription

We overlapped SVs with different categories of genes, and reported the gene profile around SVs as a number of overlapping genes per bin, per SV. For scenario (1) we worked with 100 bins per SV length, and for scenario (2), we worked with 100 bins per Mb. We defined highly expressed genes as genes whose expression was above the median. We used expression values from cancer types most represented in a given SV set.

#### APOBEC mutation positioning and strand with respect to SV breakpoints

To understand the relationship between SVs and APOBEC-attributed mutations, we sought their overlap on chromosomes. We considered only non-clustered SVs, and reported the aggregated results per SV signature. For each SV, we also sought APOBEC-attributed mutations in SV “margins”—defined as 100 kb regions on either side of the SV, irrespective of SV size. For Figures 3B, 3C we used SVs with inter-breakpoint distances lower than 300 kb and the entire spectrum of SV lengths for APOBEC strand analysis in Figure 4D.

We reported the reference strand of each APOBEC mutation overlapping with SV footprints or their margins. For mutations overlapping with the SV footprint, we calculated the relative position within the SV, dividing each SV into 10 or 2 regions. From the latter, we prepared 2×2 tables counting the mutations, with rows corresponding to the left or right side of the SV midpoint, and columns corresponding to APOBEC-attributed mutations with G or C reference alleles: 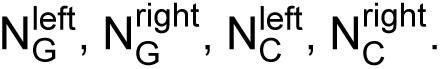We then used Fisher’s exact test to calculate the odds ratio and statistical significance. We additionally calculated a strand effect size:

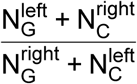

with the intuitive interpretation of the ratio of number of mutations that follow the “Gs-then-Cs” model over those that do not.

For the APOBEC analysis, we used mutations where the predicted mutation multiplicity was less than 1.5 to eliminate the APOBEC mutations that might have occurred before duplication. Mutation multiplicity estimates were calculated by MutationTimeR and retrieved from a previous publication.^49^ We verified that most of the APOBEC attributed point mutations have a predicted multiplicity of 1.5 or less; for point mutations overlapping with RS1 duplications, 76% of APOBEC attributed mutations have a multiplicity of 1.5. Eliminating the constraint on mutation multiplicity, the relationship between mutation strand and position with respect to SVs breakpoint remained highly significant (odds ratio: 3.65 [3.05–4.37], effect size 1.91, P < 1 *×* 10^-15^).

We distinguished the APOBEC mutations overlapping with the 5’ and 3’ margins. A 2×2 table summarized the APOBEC mutations overlapping with the left and right margins (rows) and the reference strand (columns) and calculated the effect size and statistical significance similarly.

#### Identifying patient subpopulations: *BRCA1*/2-/-, *CCNE1*-amplified, *CDK12*-/-

To identify PCAWG samples with bi-allelic *CDK12* loss, we used OncoKb’s variant prioritization.^50^ For conservative analysis, we identified samples with evidence of bi-allelic inactivation, which could be either because of two somatic mutations or a variant allele fraction approaching tumor purity. We identified 7 PCAWG samples with evidence of bi-allelic *CDK12* loss.

As *CCNE1*-amplified, we considered samples whose number of *CCNE1* gene copies exceeded sample ploidy by 4 copies of more.

We sought PCAWG samples with bi-allelic loss of *BRCA1* or *BRCA2* genes. As such, we considered samples where the PCAWG consortium identified pathogenic mutations, and evidence of bi-allelic loss was found in the CHORD data curation of the PCAWG dataset.^1,23^

#### New SV catalogues

SVs were first classified as clustered or not, then according to SV type and direction.^5^ We only considered non-clustered SVs for signature discovery and fitting.

- **Replication Fork Direction** As described above, OK-seq-derived replication fork direction around each SV was described as a tuple (D_1_, D_2_) where D_i_ *∈* {L, R}. Where OK-seq did not indicate a consistent replication direction, some SVs could not be classified. We then counted the SVs per sample, according to a combination of old 16 features and further 4 subclasses according to RFD.
- **Replication Timing** We used the Wavelet-smoothed replication timing weighted signal available from Encode for the MCF7 cell line. Genome-wide, the values were stratified into 5 quantiles, corresponding to “very early”,“early”,“medium”,“late” and“very late” timing for the locs during replication. SVs were then classified according to the replication timing of the SV midpoint. For inter-chromosomal translocations where midpoint cannot be defined, we used average timing of two breakpoints of an SV.

#### Discovery of SV signatures in the PCAWG dataset

The discovery of mutational signatures from non-clustered SVs categorized into 64 RFD and 80 RT-aware channels was performed using the implementation of non-negative matrix factorization in the MuSiCal package.^35^ For signature discovery, we only kept samples with at least 30 non-clustered SV each.

The PCAWG RFD extraction was performed on an SV count matrix consisting of 632 samples and 64 features (a total of 88,784 SVs). The RT extraction was performed on a count matrix of 632 samples and 80 SV features (89,037 SVs).

For RFD signature discovery, we accepted the solution with 15 signatures, as suggested by MuSiCal’s criteria that combine the stability of signatures and the potential to explain most of the data. For RT, we settled on the solution with 18 signatures, a higher number than suggested by MuSiCal. While the average silhouette score was 0.7, many of the additional and interesting signatures had silhouette scores higher than that.

#### Normalization of RFD Signatures

The direction of replication in the human genome is uneven, making the interpretation of RFD signatures challenging. When we include the RFD features in signature analysis, some types of events may be pronounced, even if no RFD configuration predisposed loci to SVs. The RFD thus needs to be corrected for the’opportunity’ of each kind of event to occur. The RFD-opportunity is likely to depend on the size and the inter-breakpoint distance of SVs. We thus simulated SVs by randomly placing them on chromosomes, matching the observed distribution of SV sizes. For each class of SVs (C: SV type x size class, e.g., 10-100 tandem duplication), we calculated the expected proportion of SVs falling into each RFD subclass (R, e.g., R-R, R-L, L-R, R-R). The correction factor f_R_^C^ was defined using the number of simulated SVs in RFD configuration R in class 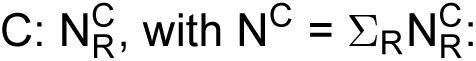

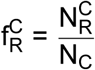

For visualizations, the signatures, e.g. 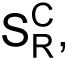 were corrected as follows:

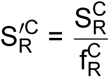

accounting for the differential availability of regions with a given RFD configuration in the genome, conditioned on SV size.

#### SVIG: multi-class classifier of sample SV patterns

The discovered RFD and RT signatures were estimated for every sample and used as features for predictions. The PCAWG training set consisted of 554 samples without a label (other), 41 samples classified as CCNE1-amplified, 18 samples with bi-allelic BRCA1 loss, 13 samples with bi-allelic BRCA2 loss, and 5 samples with bi-allelic CDK12 loss. For training, samples were weighted inversely proportional to class frequency; otherwise, the classifiers did not pay enough attention to rarer classes. The XGBClassifier, as implemented in Python’s xgboost package, was trained with a learning rate of 0.05, a maximum depth of 3, and a subsampling rate of 0.8. The out-of-fold predictions were used to calculate the suspicion values, where the model made confident predictions that were inconsistent with data labels. Suspicion values were calculated as 1 - P(C_T_), where P(C_T_) is the predicted probability of the true class per sample. Samples where this probability was lower and suspicion values were higher were downweighted. This procedure was implemented to account for the fact that more tumors may display the SV phenotypes compared to those where causal alterations are identifiable, i.e. there can be real SV phenotypes without a known cause that would appear as false positives. The decile of samples with the highest suspicion values was moderately down-weighted in training (0.8 to 1). With the amended weights, the classifier was retrained. We report confusion matrices and accuracy metrics on out-of-fold predictions, that is, when samples being predicted were not included in the training dataset. Feature importance per class was measured using Shapley values.

The PCAWG-trained model was then applied as is to the GEL dataset or, alternatively, retrained on the GEL dataset. The GEL dataset offers a considerable boost in the number of samples with confirmed HRD and replication stress phenotypes (Figure 6E).

## Data and Code Availability

We accessed the PCAWG dataset of 2,778 cancer samples. We used the consensus calls obtained from multiple single nucleotide variant callers produced by the PCAWG consortium, where evidence from at least 2 out of 4 callers (CaVEMan v1.5.1, MuTect, MuSE v1.0rc, DKFZ) was required per variant.^23^ Somatic mutation calls from the PCAWG study (.vcf files dated’20160830’) were accessed through the ICGC data portal, and we included all mutations with the’PASS’ flag. PCAWG’s consensus SV calls were downloaded from the ICGC data portal. We used the consensus SV“.bedpe” files with dated’161116’. For microhomology analysis, we used the SV calls generated by the Hartwig Medical Foundation using Gridds (v2.9.3) and Purple (v2.53) algorithms.^30,51^

Genomics England dataset. Data from the National Genomic Research Library (NGRL) used in this research are available within the secure Genomics England Research Environment. Access to NGRL data is restricted to adhere to consent requirements and protect participant privacy. We used the SV calls obtained as a consensus of multiple SV callers. We further used copy number profiles to evaluate *CCNE1* amplifications. We used mutation calls and their annotation to evaluate loss-of-function of *CDK12*. We used the calls for bi-allelic loss of HRD genes as published previously.^52^

Access to NGRL data is provided to approved researchers who are members of the Genomics England Research Network, subject to institutional access agreements and research project approval under participant-led governance. For more information on data access, visit: https://www.genomicsengland.co.uk/research

Code to reproduce the analysis in this study is available through: https://github.com/maigiclab/SVIG.

## Supporting information

Table 4: Sample labels and SVIG predictions on the PCAWG dataset.

Table 3: Redefined SV signatures with replication fork direction (RFD) and timing (RT) features, and exposures in the PCAWG set.

Table 2: APOBEC strand polarity, within and in margins of SVs, for SV signatures as defined in Degasperi et al.

Table 1: Replication, transcription, and repeat features of SV signatures as defined in Degasperi et al.

## Author Contributions

D.G. conceived the study, developed the computational methodology, performed the formal analyses, curated samples and data, managed resources, supervised the work and acquired funding. D.G. drafted the manuscript with input from all authors. T.R. contributed to conceptual development and replication-direction analyses. J.C. contributed to conceptual development of the APOBEC strand analysis. S.E. and M.A. contributed data visualizations. A.T., A.C., R.H. and D.C.W. curated structural variant data from the Genomics England (GEL) study. P.P. acquired funding. All authors reviewed and approved the final manuscript.

## Acknowledgements

M.A. and D.G. are supported by the ‘Major Grant’ award from the Fund for Innovation in Cancer Informatics awarded to D.G. and P.P.

At the Institute of Cancer Research (ICR), work was supported by Cancer Research UK (CRUK) (C1298/A25514.) and the Wellcome Trust (214388). D.C.W. is supported by the NIHR Manchester Biomedical Research Centre (NIHR203308).

We gratefully acknowledge the participants of the National Genomic Research Library (NGRL), whose contributions made this research possible. Secure access to the NGRL under project ID 1166 was provided by Genomics England, which delivers the NGRL in partnership with NHS England, and is wholly owned by the UK Department of Health and Social Care. The NGRL contains participants’ health data collected by the NHS as part of their care, along with samples and data from their participation in research, for which fully informed consent has been obtained.

This includes genomic and clinical data provided through the NHS Genomic Medicine Service, as well as data obtained through research studies, including the 100,000 Genomes Project and the Generation Study, both of which are delivered in partnership with the NHS, and from other research cohorts involving external collaborators.

We thank Michal Zimmermann, Yuwei Zhang, Yujie Guo, Xueqing Zou, and Rahul Sharma for helpful discussions and feedback on the manuscript.

## Supplemental Figures

**Figure S1.**
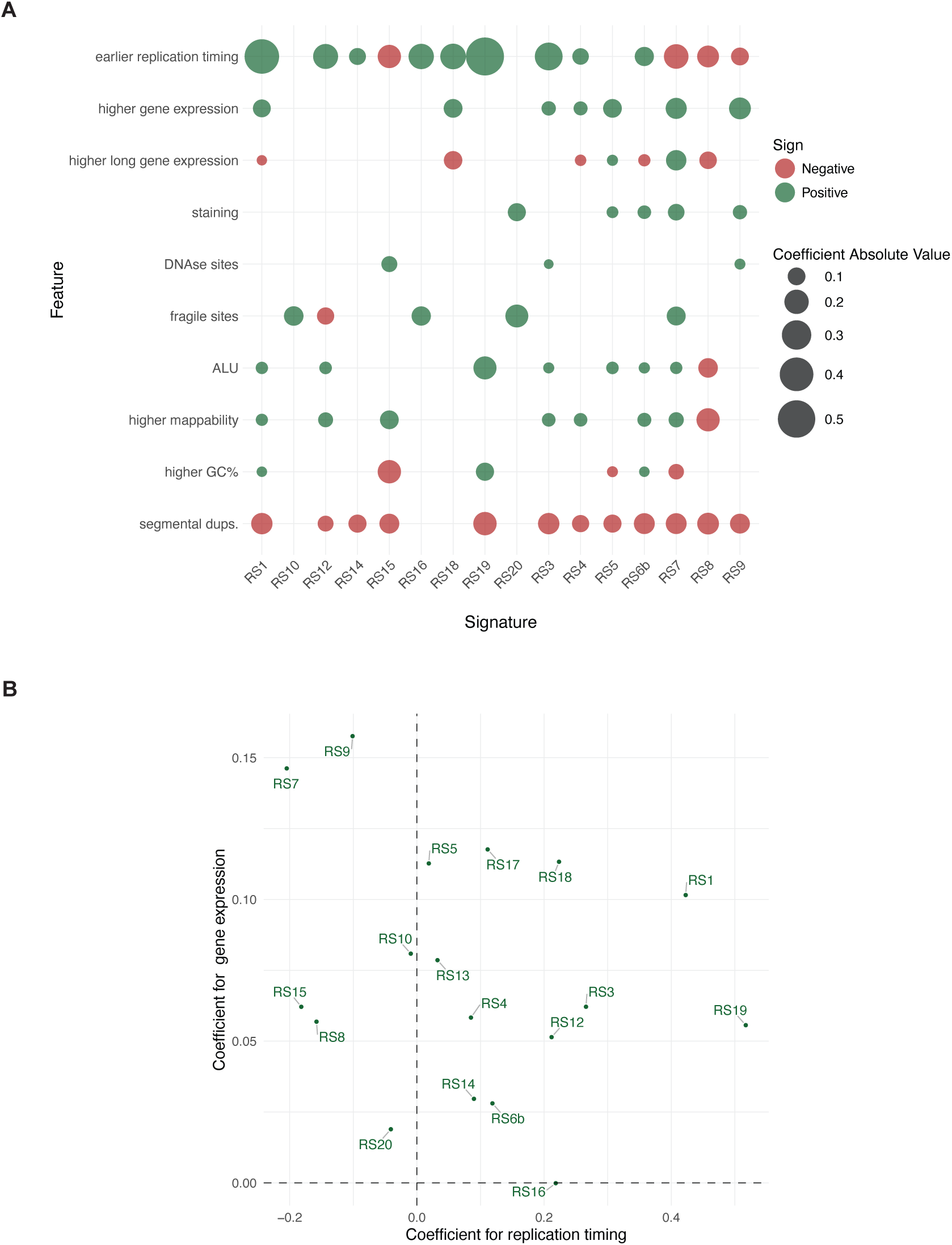
Genomic covariates of SV breakpoints analyzed at single–breakpoint resolution, treating the two breakpoints of each SV independently. A. Negative-binomial regression coefficients fitted separately for breakpoint sets defined by SV signatures. For each signature, the number of breakpoints per 1 kb bin was modeled using local genomic features as predictors. B. Scatterplot of regression coefficients for replication timing and gene expression across SV signatures (two rows from A). RS1, RS3 and RS19, duplication-dominated signatures, are most enriched in regions of early replication.

**Figure S2.**
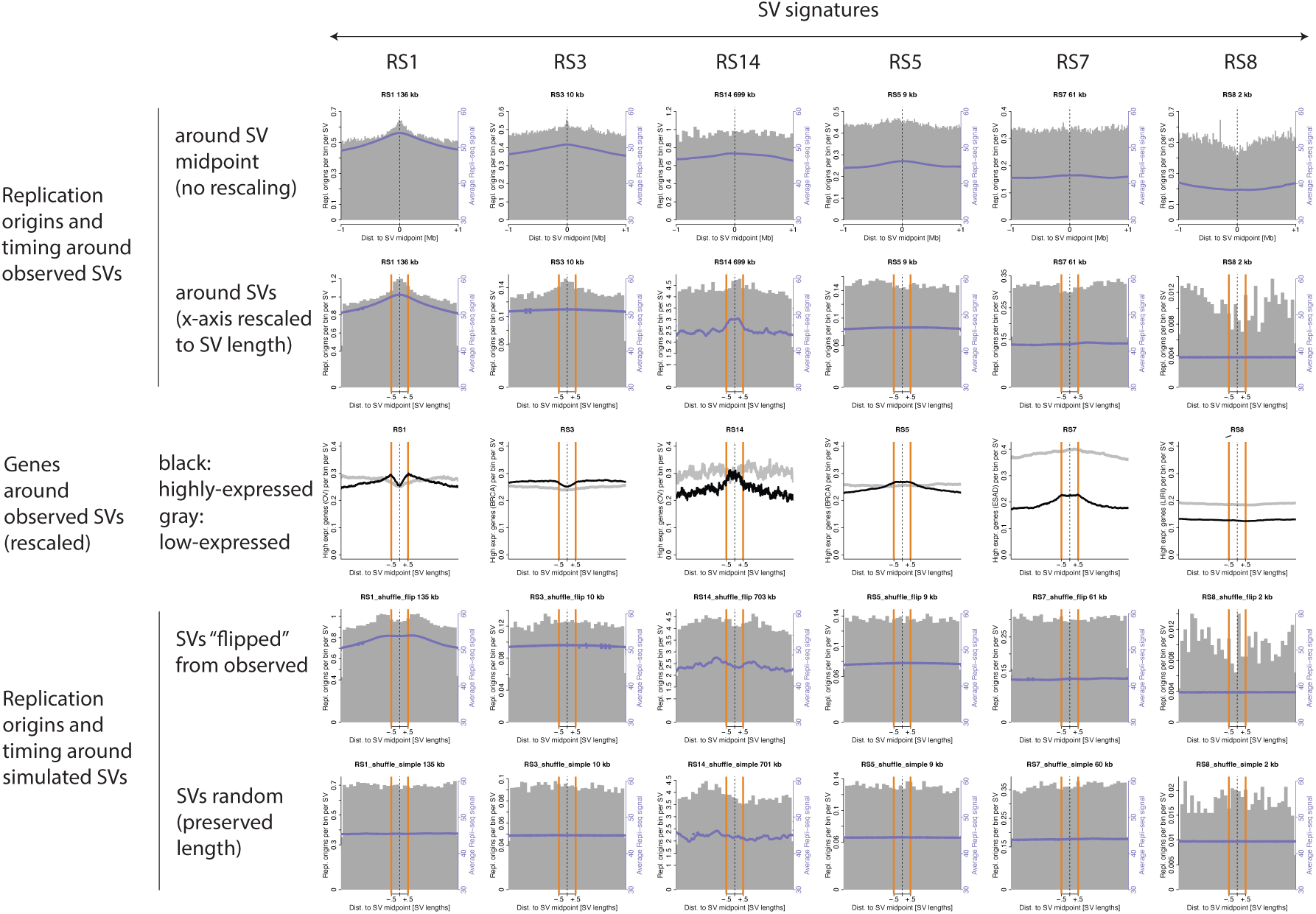
Features around SVs attributed to signatures. Columns represent SV signatures. Rows correspond to different analyses of SV positioning relative to replication origins, replication timing, and transcription. The top row shows profiles around SV midpoints without rescaling to SV length, enabling unbiased comparison between SV groups of different sizes. The bottom two rows show replication timing and origin analyses around randomly permuted SVs.

**Figure S3.**
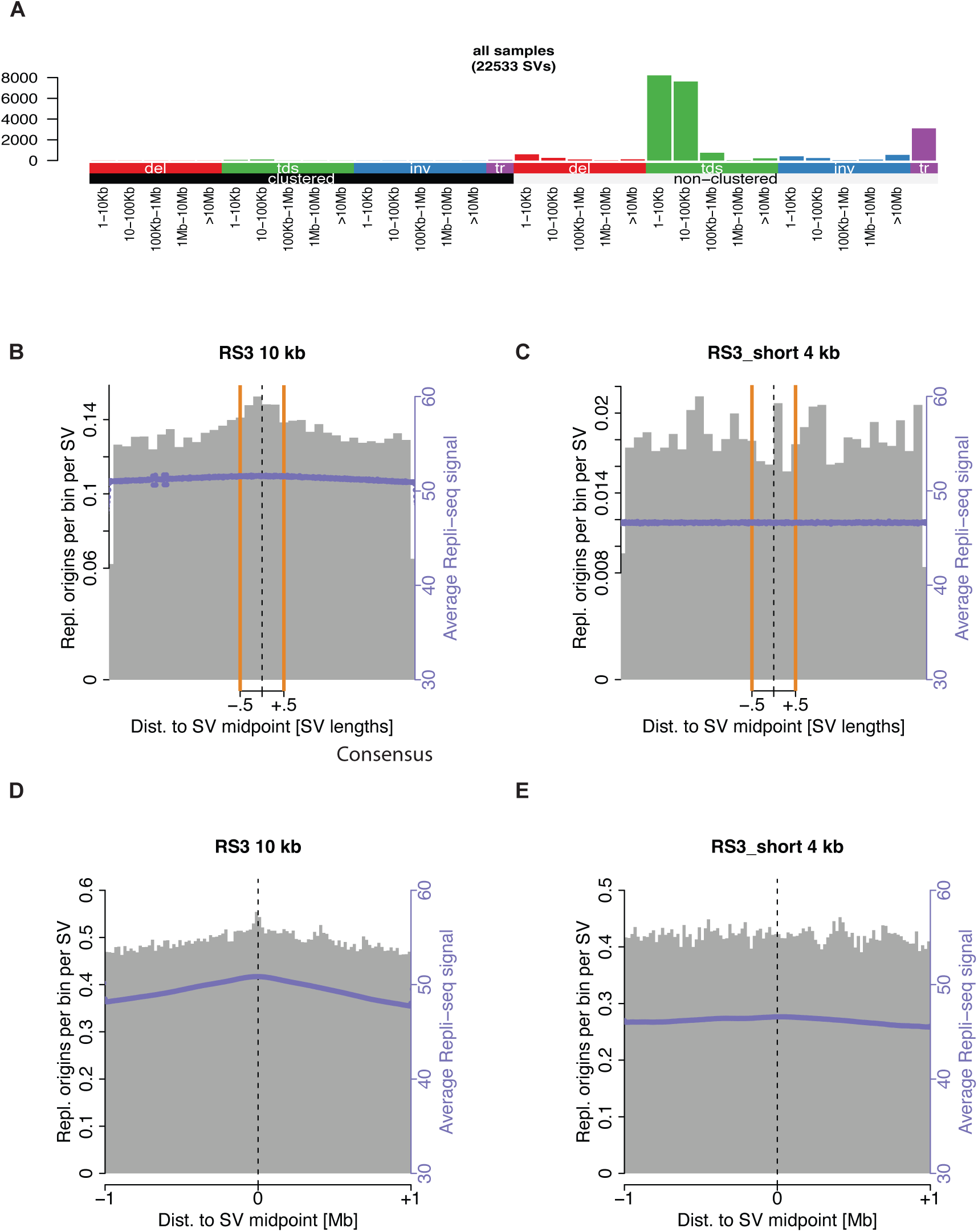
Replication origin and time profiles around RS3 SVs. 1–100 kb tandem duplications constitute the majority of RS3-attributed SVs. A shows the profile of all RS3-attributed SVs, highlighting predominantly non-clustered duplications of length 1–100 kb. B and D show replication timing and replication-origin profiles around all RS3-attributed SVs, with SVs rescaled to unit length in B and shown at their native lengths in D. C (rescaled) and E (non-rescaled) show RS3-attributed SVs shorter than 10 kb. D and E are directly comparable, demonstrating that the excess of replication origins at RS3 midpoints is restricted to SVs longer than 10 kb.

**Figure S4.**
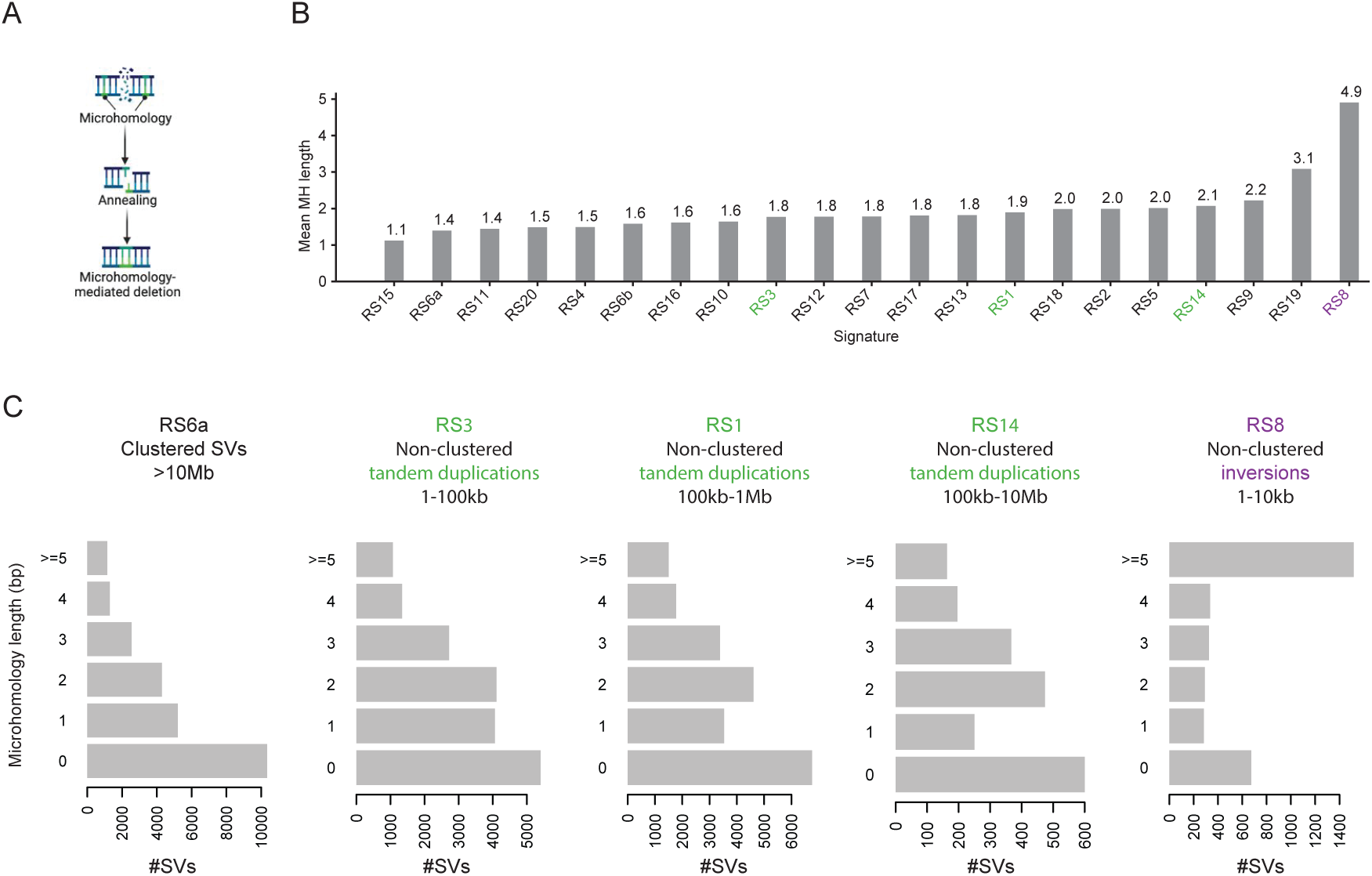
Microhomology at breakpoints of SVs attributed to mutational signatures. A: Schematic of microhomology at SV breakpoints. B: Mean microhomology length at SV breakpoints across signatures. Chromothripsis-associated signatures (RS6a, RS6b) show among the lowest values, whereas the short (<10 kb) inversion signature RS8 shows the highest. C: SV signature catalogues stratified by microhomology length. For tandem duplication signatures (RS1, RS3, RS14), mean microhomology increases with SV size, while short foldback inversions attributed to RS8 show an outlier distribution.

**Figure S5.**
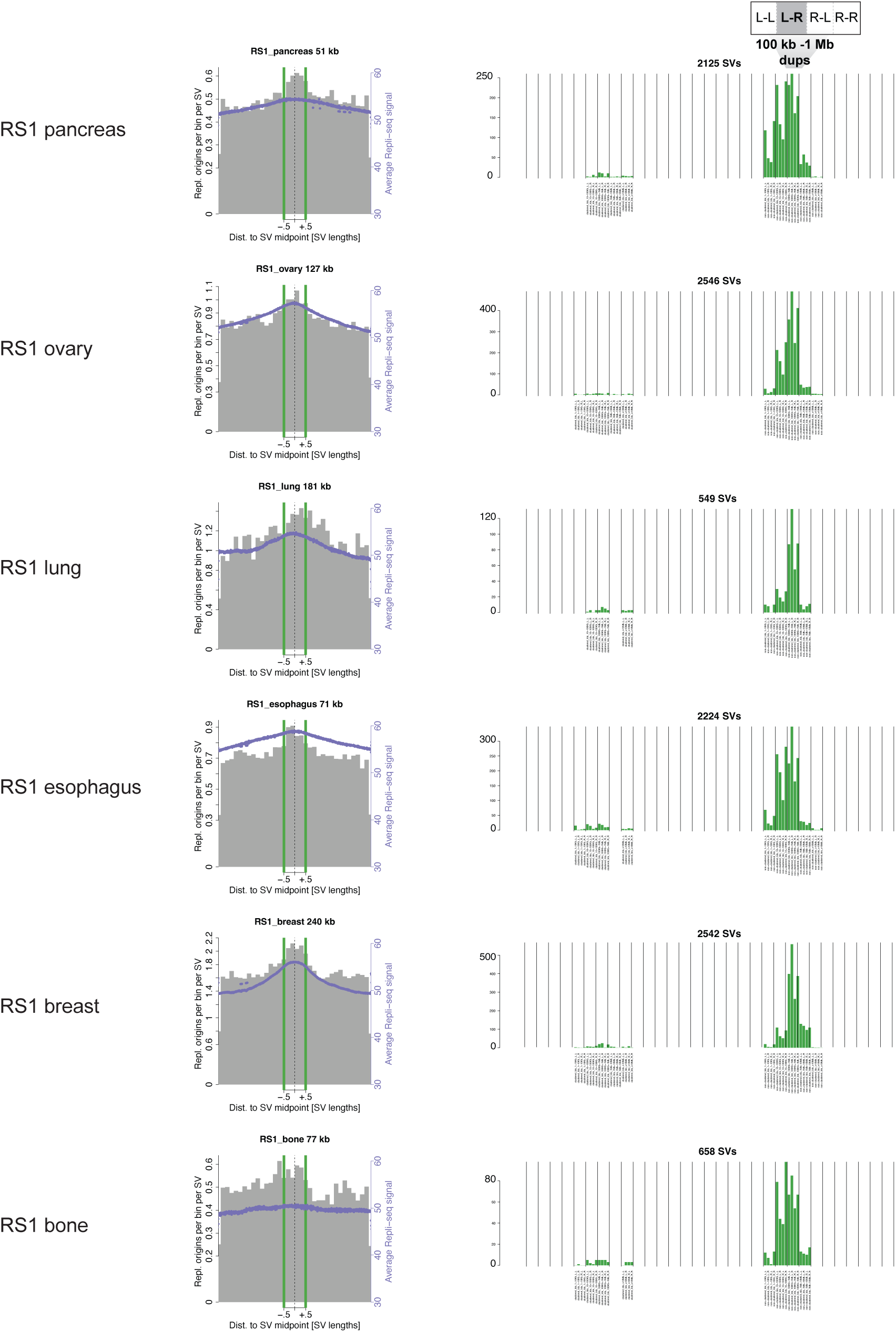
Replication origin and fork-direction features of RS1-attributed SVs across tissues. Rows correspond to RS1 analyses across different tissues. The left column shows mean replication timing and replication-origin density around RS1-attributed tandem duplications (predominantly 100 kb–1 Mb, nonclustered), after rescaling each SV to unit length (median SV size shown above). The right column summarizes RS1- attributed tandem duplications stratified by replication-fork direction at the two breakpoints (L–L, L–R, R–L, R–R). SV counts are not normalized for the genomic distribution of replication-fork direction. In size-matched simulations, the L–R configuration is less frequent than the L–L and R–R configurations. In contrast, in the observed data, the L–R orientation predominates among 100 kb–1 Mb events across all tissues.

**Figure S6.**
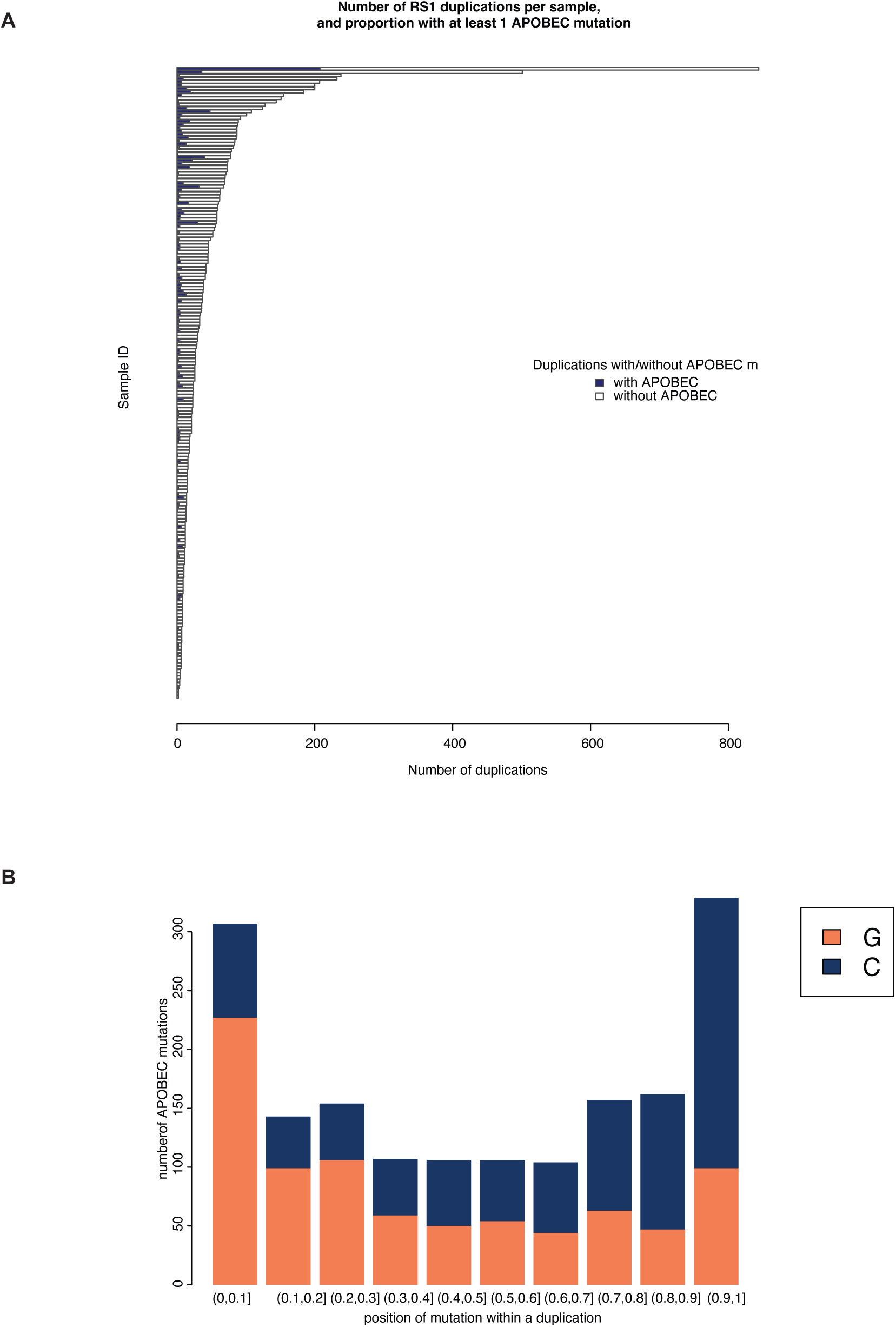
Coincidence of APOBEC-mutations with RS1-attributed SVs. A. Sample distribution of RS1-attributed SVs that overlap with APOBEC-attributed point mutations; B. Nonnormalized distribution (counts) of APOBEC mutations overlapping with RS1-attributed SVs.

**Figure S7.**
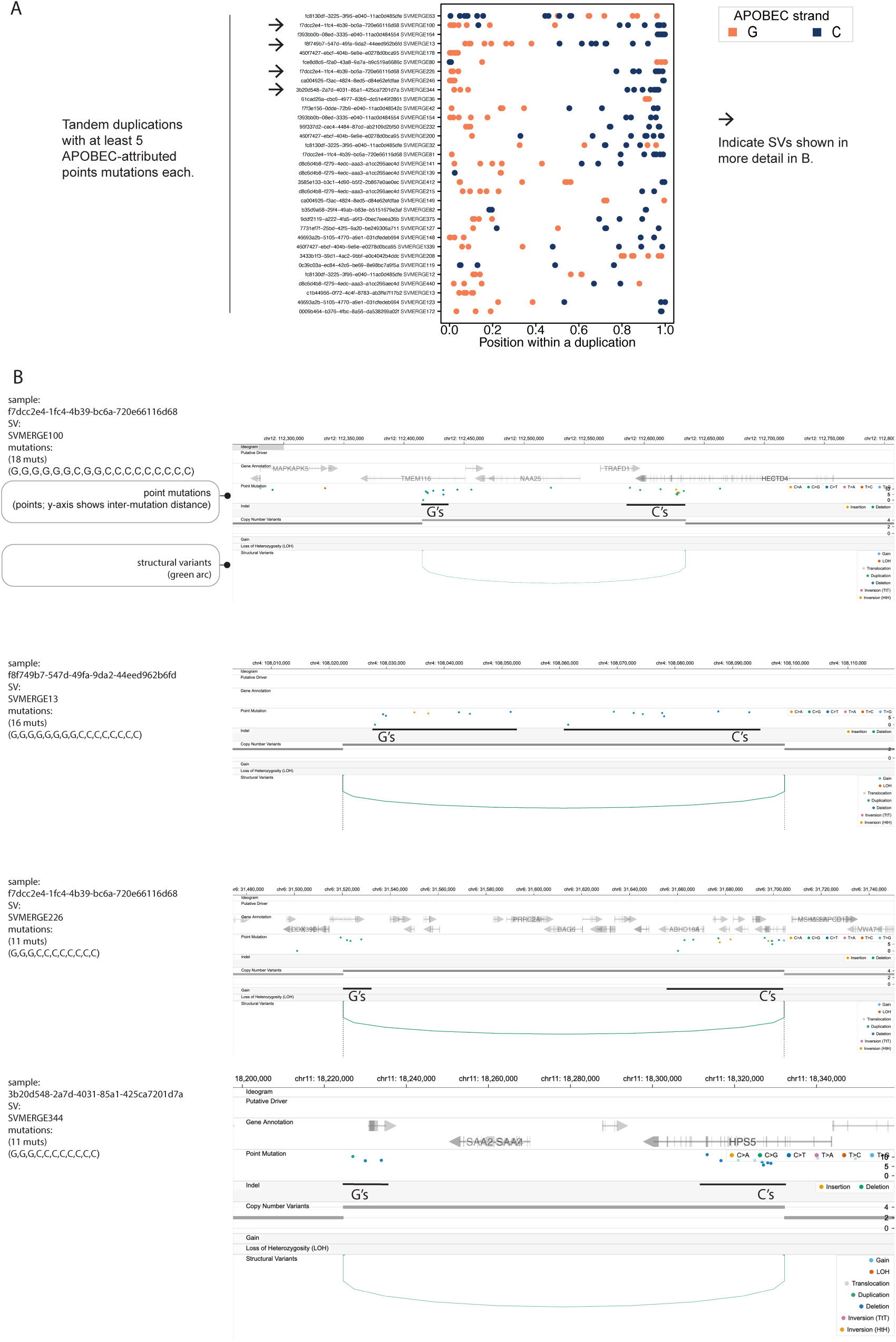
Visualizations of selected tandem duplications and adjacent strand of APOBEC-attributed mutations. A. Selection of tandem duplications and adjacent APOBEC-attributed mutations exhibiting the dominant pattern of G>N mutations near the left breakpoint and C>N mutations near the right breakpoint. The plot is limited to SVs containing at least five such point mutations. Arrows indicate SVs shown in more detail in B. B. Chromoscope.bio-generated visualizations of selected SVs, showing tandem duplications and point mutations. Tandem duplications are shown as green arches, and only APOBEC-attributed mutations are displayed, highlighting regions with consistent G calls on the negative strand and C calls on the positive strand. Only duplications with at least five APOBEC-attributed mutations are shown. The APOBEC-attributed mutations follow the G-then-C pattern and occur close to SV breakp_8_oints.

**Figure S8.**
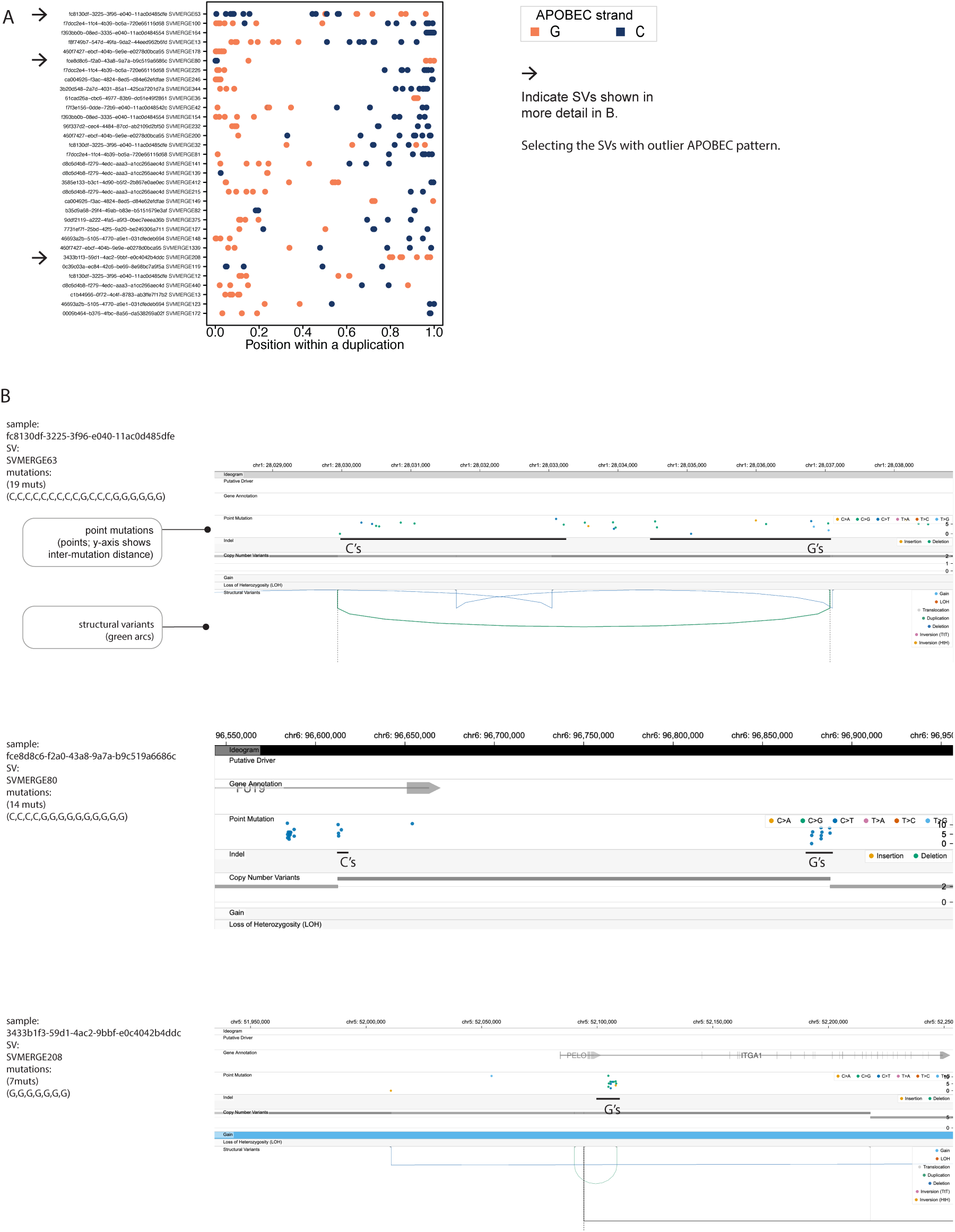
APOBEC-attributed mutations around SVs exhibit an outlier strand pattern. A. Duplications with APOBEC-attributed mutations show the inverse of the dominant pattern, with C>N mutations near the left breakpoint and G>N mutations near the right breakpoint. B. Chromoscope.bio-generated visualizations. Two of the three events are complex SVs; when a duplication represents a secondary event, the rearranged chromosome may exhibit this atypical APOBEC strand pattern.

**Figure S9.**
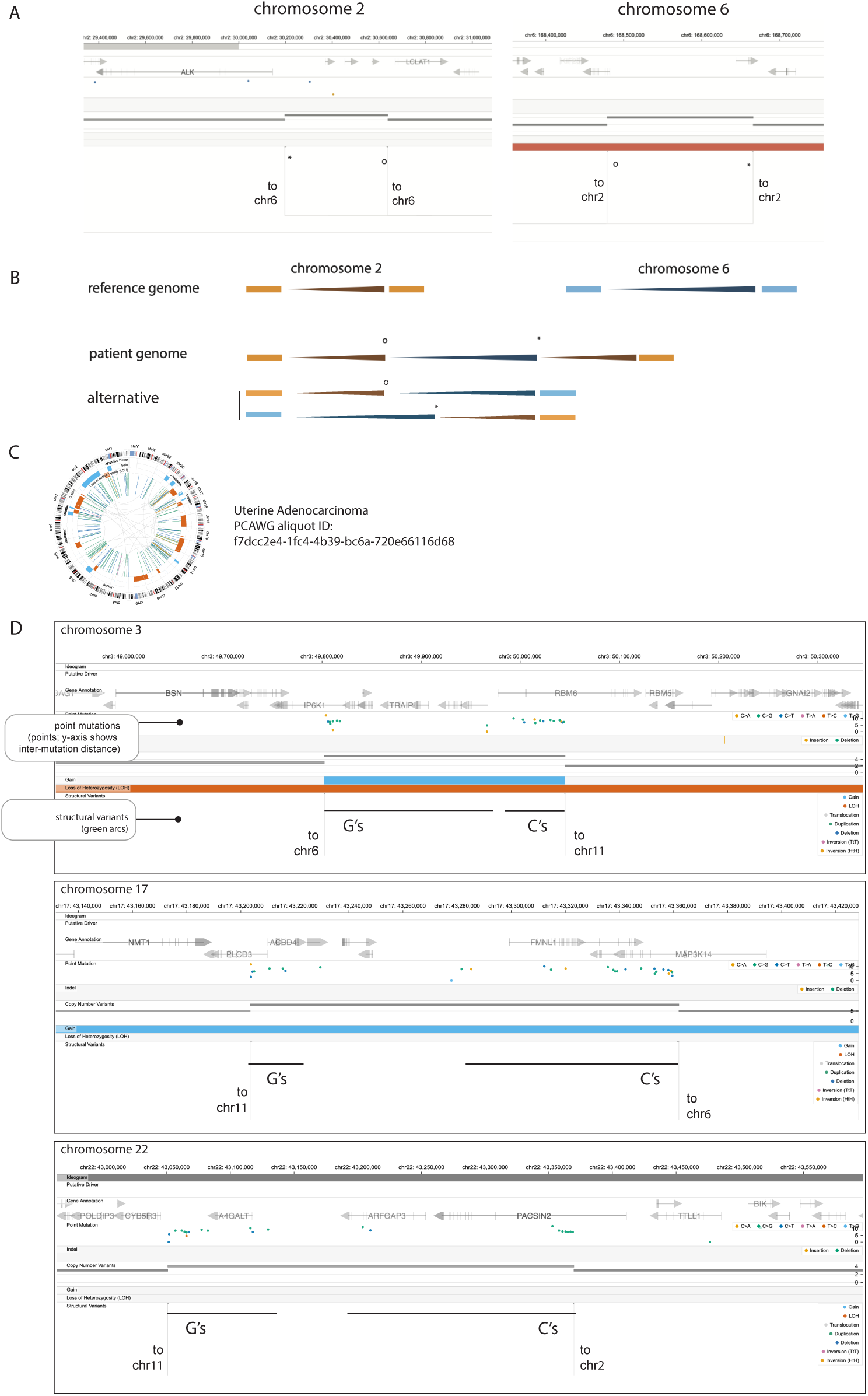
Chromosomal gains defined by semi-balanced translocations exhibit a ‘Gs-then-Cs’ APOBEC strand pattern. A–B. Diagram of a chromosomal gain delineated by a semi-reciprocal translocation, also referred to as a ‘synthetic duplication’ or ‘templated insertion’. C. Example uterine cancer genome with a tandem duplicator phenotype and high APOBEC mutagenesis. D. APOBEC-attributed mutations within the gained regions follow the same polarity pattern observed in tandem duplications.

**Figure S10.**
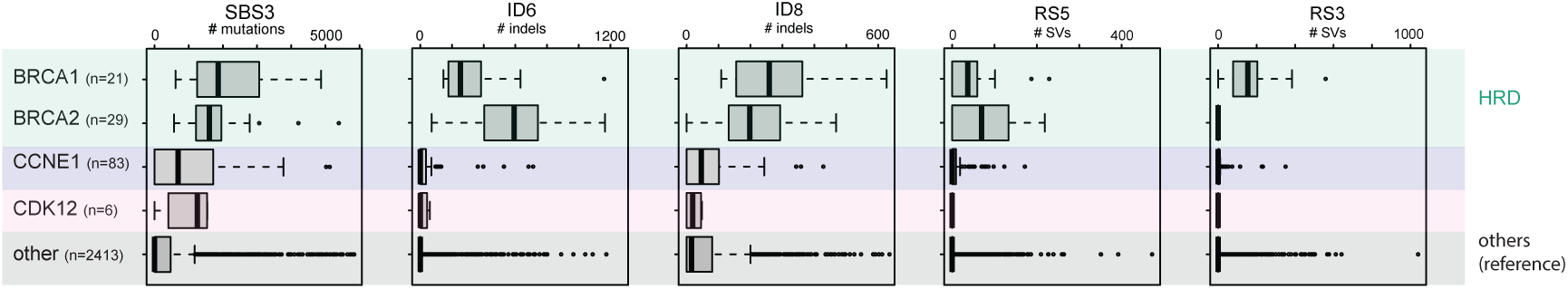
Comparison of point mutation– and indel-based mutational signature burdens stratified by gene alterations. *CDK12*-mutant and *CCNE1*-amplified tumors show SBS3 but do not exhibit the indel (ID6, ID8) or structural-variant (RS3, RS5) signatures typically associated with homologous recombination deficiency.

**Figure S11.**
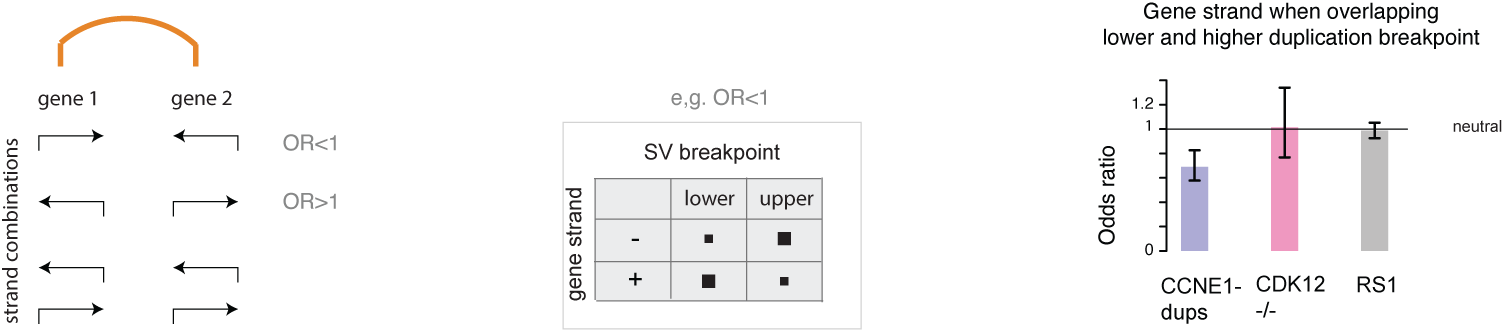
Association between gene transcriptional strand and the SV breakpoint overlapping the gene. *CCNE1*-amplified tumors exhibit transcriptional strand bias, with positive-strand genes preferentially overlapping lower tandem duplication breakpoints and negative-strand genes overlapping upper breakpoints (Fisher’s exact test; 95% confidence intervals).

**Figure S12.**
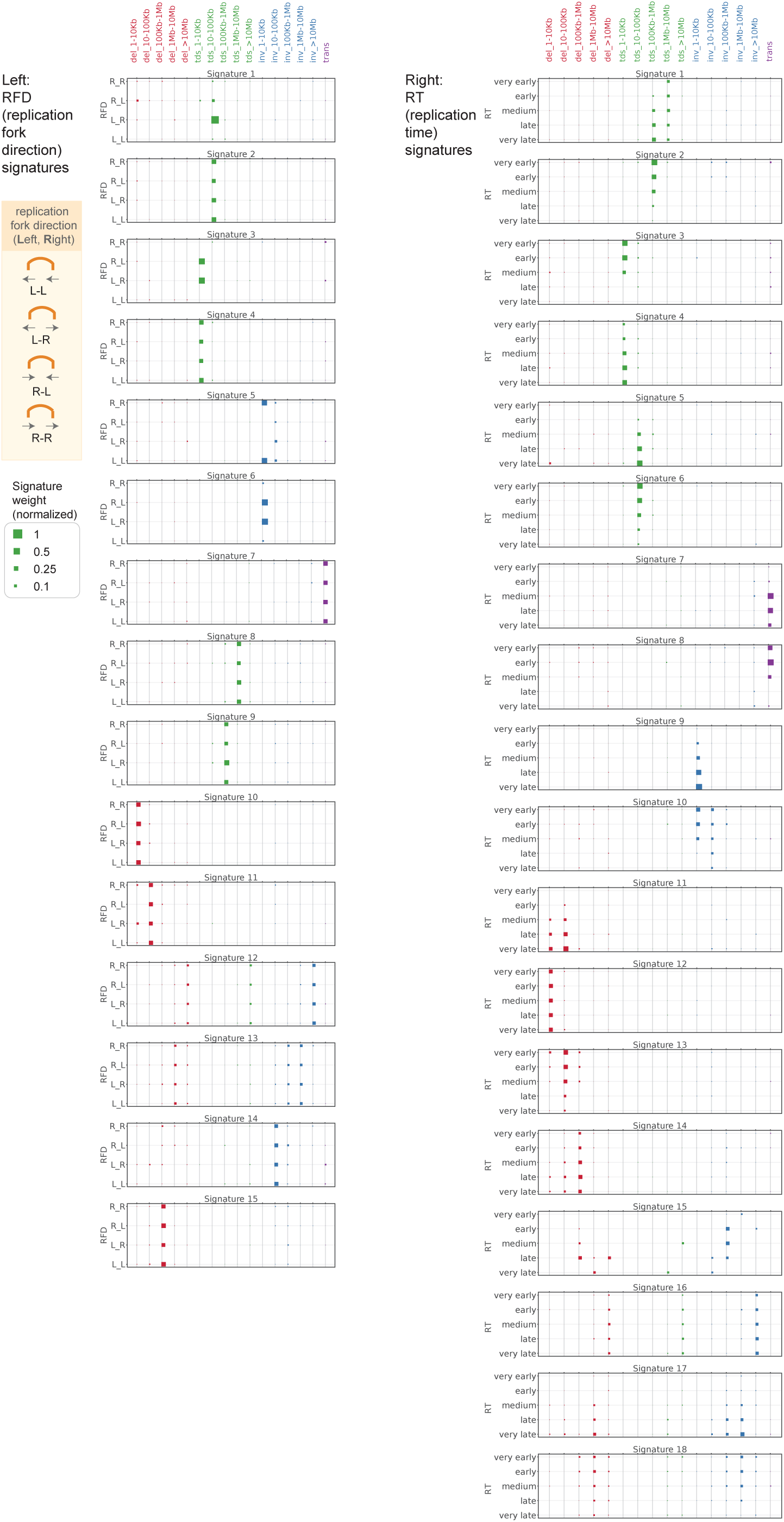
SV signatures derived from PCAWG data. Left: replication fork direction (RFD) signatures. Right: replication timing (RT) signatures. Signatures were extracted from non-clustered SVs across PCAWG cohorts (breast, ovary, prostate, uterus, and upper gas_1_t_2_rointestinal). Incorporating replication features resolves several previously defined signatures into two signatures, exemplified by pairs of RFD signatures 3 and 4 and RT signatures 3 and 4.

**Figure S13.**
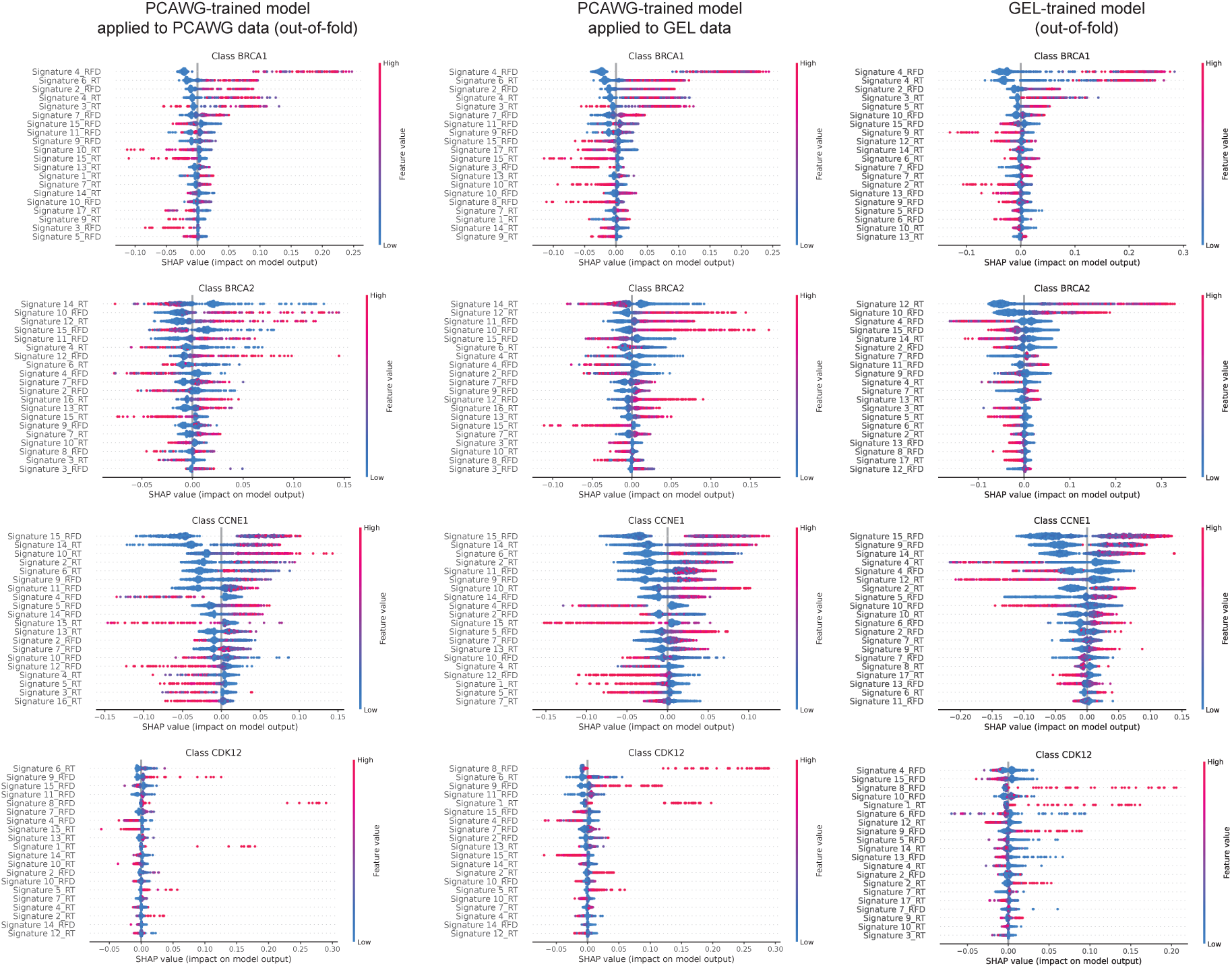
Feature importance for class predictions made by SVIG. Feature importance is quantified using Shapley values. Left: Pan-Cancer Analysis of Whole Genomes (PCAWG) data evaluated with the PCAWG-trained model. Middle: Genomics England (GEL) data evaluated with the PCAWG-trained model. Right: GEL data evaluated with the GEL-retrained model. Multiple signatures contribute to class predictions. Replication timing–defined features influence predictions: RT6 is a positive predictor of the *CCNE1* class, whereas RT5 is a negative predictor, despite both being predominantly composed of 10–100 kb tandem duplications.

